# Maturation, developmental site, and pathology dictate murine neutrophil function

**DOI:** 10.1101/2021.07.21.453108

**Authors:** John B. G. Mackey, Amanda J. McFarlane, Thomas Jamieson, Rene Jackstadt, Ximena L. Raffo-Iraolagoitia, Judith Secklehner, Xabier Cortes-Lavaud, Frédéric Fercoq, William Clarke, Ann Hedley, Kathryn Gilroy, Sergio Lilla, Juho Vuononvirta, Gerard J. Graham, Katia De Filippo, Daniel J. Murphy, Colin W. Steele, Jim C. Norman, Thomas G. Bird, Derek A. Mann, Jennifer P. Morton, Sara Zanivan, Owen J. Sansom, Leo M. Carlin

## Abstract

Neutrophils have been implicated in poor outcomes in cancer and severe inflammation. We found that neutrophils expressing intermediate levels of Ly6G (Ly6G^Int^) were present in mouse cancer models and more abundant in those with high rates of spontaneous metastasis. Maturation, age, tissue localization and functional capacity all drive neutrophil heterogeneity. Recent studies have proposed various markers to distinguish between these heterogeneous sub-populations; however, these markers are limited to specific models of inflammation and cancer. Here, we identify and define Ly6G expression level as a robust and reliable marker to distinguish neutrophils at different stages of maturation. Ly6G^Int^ neutrophils were bona fide ‘immature neutrophils’ with reduced immune regulatory and adhesion capacity. Whereas the bone marrow is a more recognised site of granulopoiesis, the spleen also produces neutrophils in homeostasis and cancer. Strikingly, neutrophils matured faster in the spleen than in the bone marrow with unique transcriptional profiles. We propose that developmental origin is critical in neutrophil identity and postulate that neutrophils that develop in the spleen supplement the bone marrow by providing an intermediate more mature reserve before emergency haematopoiesis.

## Introduction

Neutrophils comprise a complex heterogeneous myeloid subset that becomes increasingly diverse with disease progression^1^. Neutrophil heterogeneity has been proposed to be influenced by a multitude of factors including maturation stage and age^2^, tissue localization^3^ and functional capacity^4^. Importantly, there is significant overlap between these heterogeneous neutrophil phenotypes. For example, immature neutrophils in the spleen are less motile, respond differently to bacterial infection, and localize differently when compared to mature splenic neutrophils^5, 6^.

Stages of neutrophil maturation can be characterised by distinct morphological and transcriptional phenotypes that are closely associated with functional capacity^7–9^. In disease, immature neutrophils can be prematurely released from the bone marrow (BM) and have been identified in a range of pathologies including cancer^4, 10^, sepsis^11^, and more recently severe SARS-CoV-2 infection^12^, where they have been proposed as biomarkers for disease progression and outcome. Differences in the functional capacity of immature and mature neutrophils have been proposed in mice and humans, including NETosis^13^, metabolism^14^, chemotaxis and oxidative capacity^10^. However, diversity in the methods used to identify immature neutrophils combined with the range of models investigated makes comprehensive interpretation of results in this area challenging^15^. Recent evidence for tissue specific transcriptional phenotypes^3^ and disease induced transcriptional and developmental alterations^8, 16^ further add to this complexity.

The functional influence and disease importance of neutrophil development and maturation remain poorly understood. Here, we provide comprehensive evidence that the maturation landscape and developmental niche are crucial to our understanding of the role of neutrophils in disease. Neutrophils expressing intermediate levels of (Ly6G^Int^) were identified in several mouse cancer models. These were demonstrably less mature than their Ly6G-high (Ly6G^Hi^) counterparts with specific transcriptional profiles and reduced functional capacity. Identification of maturation status based on Ly6G expression was accurate across naïve, acute inflammatory and cancer models where in the latter Ly6G^Int^ neutrophil abundance associated with metastatic potential. Furthermore, we identified that the site of neutrophil development influences their maturation and phenotype. Although the BM is a more recognised site of granulopoiesis, the spleen also produces neutrophils in homeostasis and cancer. We found that immature splenic-derived neutrophils underwent a more rapid maturation with altered transcriptomic, phenotypic, and functional capacity in a steady state and in cancer models compared to their BM-derived counterparts. We propose that this indicates a distinct role for splenic-derived neutrophils in innate immunity as a secondary source of more functionally competent / active neutrophils that supplement circulating and Ly6G^Hi^ BM reserve cells.

## Results

### Ly6G^Int^ neutrophil abundance associates with metastatic potential in mouse cancer models

We studied the neutrophils from a panel of preclinical mouse cancer models. In homeostasis, peripheral blood neutrophils are almost exclusively Ly6G^Hi^, with neutrophils expressing intermediate Ly6G levels (Ly6G^Int^) restricted to the BM (**Fig. 1a, full gating strategy Extended Data Fig. 1a**). Interestingly, we observed shifts in Ly6G expression in several of the cancer models studied. Ly6G^Int^ neutrophils in the peripheral blood and tissues of metastatic colorectal cancer (CRC) *Villin-cre*^ER^; *<u>K</u>ras*^G12D/+^; *Tr<u>p</u>53*^fl/fl^; *Rosa26*<u>^N^</u>^1icd/+^ (KPN) mice^17^ increased with tumour progression, reaching significance at clinical endpoint compared to day 30 post-tamoxifen induction controls (**Fig. 1b-d**). In contrast, Ly6G^Int^ neutrophils were not increased in the poorly metastatic models tested. These were lung adeno*-Cre* driven *<u>K</u>Ras*^G12D/+^*; Rosa26^lsl-<u>M</u>yc/lsl-Myc^* (KM)^18^ lung adenocarcinoma bearing mice induced at a virus titre where metastases do not form before endpoint (**Fig. 1e-g**), and Diethylnitrosamine-American lifestyle induced obesity syndrome (DEN/ALIOS) driven hepatocellular carcinoma bearing mice (**Fig. 1h-j**). To determine whether Ly6G^Int^ neutrophils associate with metastatic potential in the same cancer type, we compared poorly-metastatic *<u>K</u>ras*^G12D/+^*; Tr<u>p</u>53*^fl/+^*; Pdx1-<u>C</u>re* (KP^fl^C) and highly-metastatic *<u>K</u>ras*^G12D/+^*; Tr<u>p</u>53*^R172H/+^; *Pdx1-<u>C</u>re* (KPC) mouse models of pancreatic ductal adenocarcinoma (PDAC)^19^ to age-matched wild-type (WT) and *<u>K</u>ras*^G12D/+^; *Pdx1-<u>C</u>re* (KC; where only pancreatic intraepithelial neoplasia develops) controls (**Fig. 1k**). Here, KP^fl^C and KPC mice were taken with a similarly sized palpable primary tumour but prior to overt metastasis. Ly6G^Int^ neutrophils increased systemically from WT to KC, KP^fl^C and KPC respectively, correlating with the presence of lesions and metastatic potential (**Fig. 1l-p**). This increase was most apparent in the lung and liver of KPC mice, the two primary sites of metastasis in this model (**Fig. 1o, p**). Thus, the abundance of Ly6G^Int^ neutrophils mirrored metastatic potential in a panel of mouse models of cancer with different origins and drivers.

**Fig 1.**
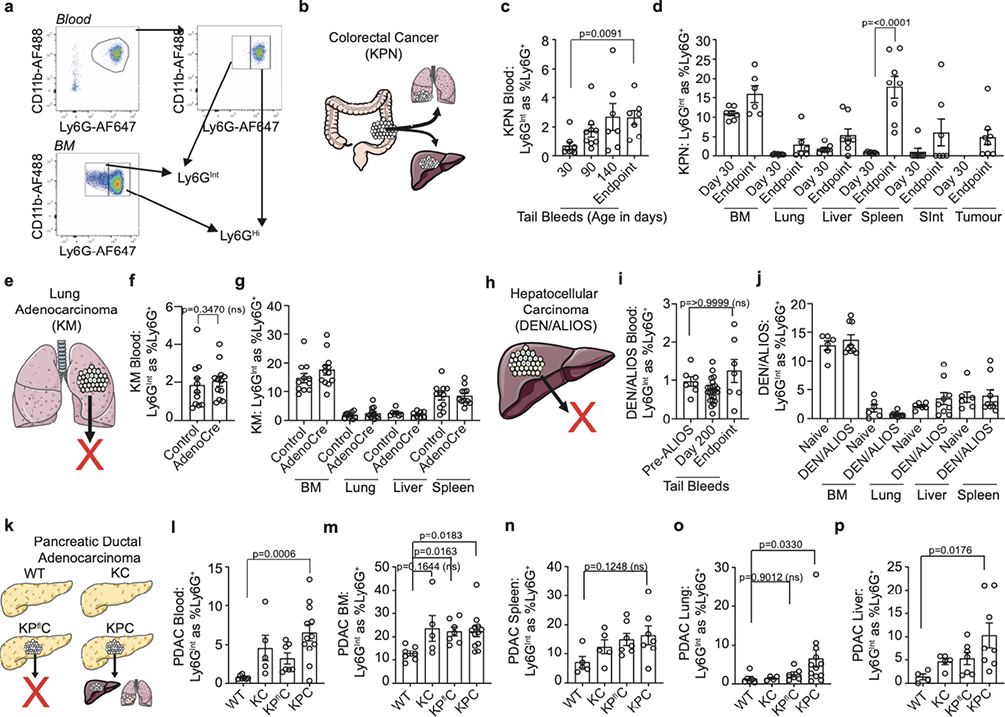
Ly6G^Int^ neutrophils associate with metastasis and metastatic potential. **a**, Ly6G^Int^ and Ly6G^Hi^ neutrophil populations were identified from the Ly6G^+^CD11b^+^ population. Ly6G^Hi^ neutrophils are predominant in the peripheral blood of naïve mice and were gated. All Ly6G^+^ neutrophils not falling into the Ly6G^Hi^ population were gated as Ly6G^Int^. **b**, Schematic for metastatic colorectal cancer mouse model, termed KPN. **c**, Quantification of Ly6G^Int^ as %Ly6G^+^ neutrophils in the peripheral blood of KPN mice. Day 30 n=8 mice; day 90 n=9 mice; day 140 n=7 mice; endpoint n=7 mice. Data were analysed by Kruskal-Wallis test with Dunn’s multiple comparisons test. **d**, Quantification of Ly6G^Int^ as %Ly6G^+^ neutrophils in the tissues of KPN mice. Bone marrow (BM): day 30 n=7 mice; endpoint n=6 mice. Lung: day 30 n=7 mice; endpoint n=6 mice. Liver: day 30 n=7 mice; endpoint n=8 mice. Spleen: day 30 n=7 mice; endpoint n=8 mice. Small intestine (SInt): day 30 n=7 mice; endpoint n=8 mice. Tumour: endpoint n=8 mice. Data were analysed by 2way ANOVA with Sidak’s multiple comparisons test. **e**, Schematic for non-metastatic lung adenocarcinoma mouse model, termed KM. **f**, Quantification of Ly6G^Int^ as %Ly6G^+^ neutrophils in the peripheral blood of KM mice. Control n=11 mice; AdenoCre n=12 mice. Data were analysed by Mann-Whitney test. **g**, Quantification of Ly6G^Int^ as %Ly6G^+^ neutrophils in the tissues of KM mice. BM: control n=11 mice; AdenoCre n=13 mice. Lung: control n=11 mice; AdenoCre n=12 mice. Liver: control n=7 mice; AdenoCre n=8 mice. Spleen: control n=11 mice; AdenoCre n=12 mice. Data were analysed by 2way ANOVA with Sidak’s multiple comparisons test. **h**, Schematic for non-metastatic hepatocellular carcinoma mouse model, termed DEN/ALIOS. **i**, Quantification of Ly6G^Int^ as %Ly6G^+^ neutrophils in the peripheral blood of DEN/ALIOS mice. Pre-ALIOS n=7 mice; day 200 n=23 mice; endpoint n=7 mice. Data were analysed by Kruskal-Wallis test with Dunn’s multiple comparisons test. **j**, Quantification of Ly6G^Int^ as %Ly6G^+^ neutrophils in the tissues of DEN/ALIOS mice. All tissues: naïve n=6 mice; DEN/ALIOS n=9 mice. Data were analysed by 2way ANOVA with Sidak’s multiple comparisons test. **k**, Schematic for WT-control, KC-control, KP^fl^C non-metastatic and KPC metastatic mouse models of pancreatic ductal adenocarcinoma. **l**, Quantification of Ly6G^Int^ as %Ly6G^+^ neutrophils in the peripheral blood of WT, KC, KP^fl^C and KPC mice. WT n=7 mice; KC n=5 mice; KP^fl^C n=7 mice; KPC n=12 mice. Data were analysed by Kruskal-Wallis test with Dunn’s multiple comparisons test. **m**, Quantification of Ly6G^Int^ as %Ly6G^+^ neutrophils in the BM of WT, KC, KP^fl^C and KPC mice. WT n=7 mice; KC n=5 mice; KP^fl^C n=7 mice; KPC n=12 mice. Data were analysed by Kruskal-Wallis test with Dunn’s multiple comparisons test. **n**, Quantification of Ly6G^Int^ as %Ly6G^+^ neutrophils in the spleen of WT, KC, KP^fl^C and KPC mice. WT n=6 mice; KC n=4 mice; KP^fl^C n=6 mice; KPC n=12 mice. Data were analysed by Kruskal-Wallis test with Dunn’s multiple comparisons test. **o**, Quantification of Ly6G^Int^ as %Ly6G^+^ neutrophils in the lung of WT, KC, KP^fl^C and KPC mice. WT n=4 mice; KC n=5 mice; KP^fl^C n=7 mice; KPC n=8 mice. Data were analysed by Kruskal-Wallis test with Dunn’s multiple comparisons test. **p**, Quantification of Ly6G^Int^ as %Ly6G^+^ neutrophils in the liver of WT, KC, KP^fl^C and KPC mice. WT n=5 mice; KC n=5 mice; KP^fl^C n=7 mice; KPC n=7 mice. Data were analysed by Kruskal-Wallis test with Dunn’s multiple comparisons test. Dots in Fig. 1c-d, f-g, i-j, l-p represent individual mice. Error bars in Fig. 1c-d, f, g, i, j, l-p represent mean±SEM.

### Ly6G surface expression accurately identifies neutrophil maturity

Neutrophils that express intermediate levels of Ly6G are thought to be immature, but this hasn’t been comprehensively tested and compared with other signifiers of maturity. Therefore, we next sought to test the use of Ly6G expression as a universal marker for neutrophil maturity. Consistent with a more immature population, Ly6G^Int^ neutrophils were: restricted to the BM in a steady state, had reduced side-scatter by flow cytometry indicative of lower granularity, had a less segmented nuclear morphology compared to Ly6G^Hi^ neutrophils, and constituted a greater percentage of the Ly6G^+^CD11b^+^ populations in the BM and peripheral blood of CSF-3- and LPS-challenged mice (**Fig. 2a-c** **and Extended Data Fig. 2a, b**). Furthermore, when we cultured BM-derived Ly6G^Int^ neutrophils, they increased cell surface Ly6G and were more viable over 20 hours than their Ly6G^Hi^ counterparts (**Fig. 2d-f**). Neutrophils exit the mitotic pool during transition from the myelocyte to metamyelocyte stages of development^20, 21^. Cells that incorporated the thymidine analogue EdU during mitosis composed a greater percentage of Ly6G^Int^ compared to Ly6G^Hi^ neutrophils in the BM of PBS-control mice between 4 and 24 hours post-EdU incorporation (**Fig. 2g**). These results were replicated in G-CSF (CSF-3)-^22, 23^ and LPS-challenge^24^ mouse models to induce haematopoietic stress in which immature neutrophils are identified outside of their medullary niche in mice and humans.(**Fig. 2g**). In contrast to PBS controls, EdU^+^ neutrophils were identified in the peripheral blood as early as 24 hours post-EdU injection in CSF-3- and LPS-challenged mice (**Extended Data Fig. 2c**). Of these, a significantly higher percentage of Ly6G^Int^ neutrophils were EdU^+^ compared to the Ly6G^Hi^ population (**Fig. 2g**). Ki-67 staining confirmed that a proportion of BM Ly6G^Int^ neutrophils were outside of G0, however, Ki-67 expression was drastically reduced in the BM of CSF-3- and LPS-challenged mice (**Extended Data Fig. 2d**). These data are consistent with Ly6G^Int^ neutrophils being more recently actively mitotic compared to Ly6G^Hi^ neutrophils and a shift from cell cycle regulation to quicker post-mitotic maturation and efflux from the BM in states of haematopoietic stress^16^. Transcriptomic analysis showed clear differences between Ly6G^Int^ and Ly6G^Hi^ neutrophils dependent on tissue and stimulus (**Extended Data Fig. 2e, f**).

**Fig. 2.**
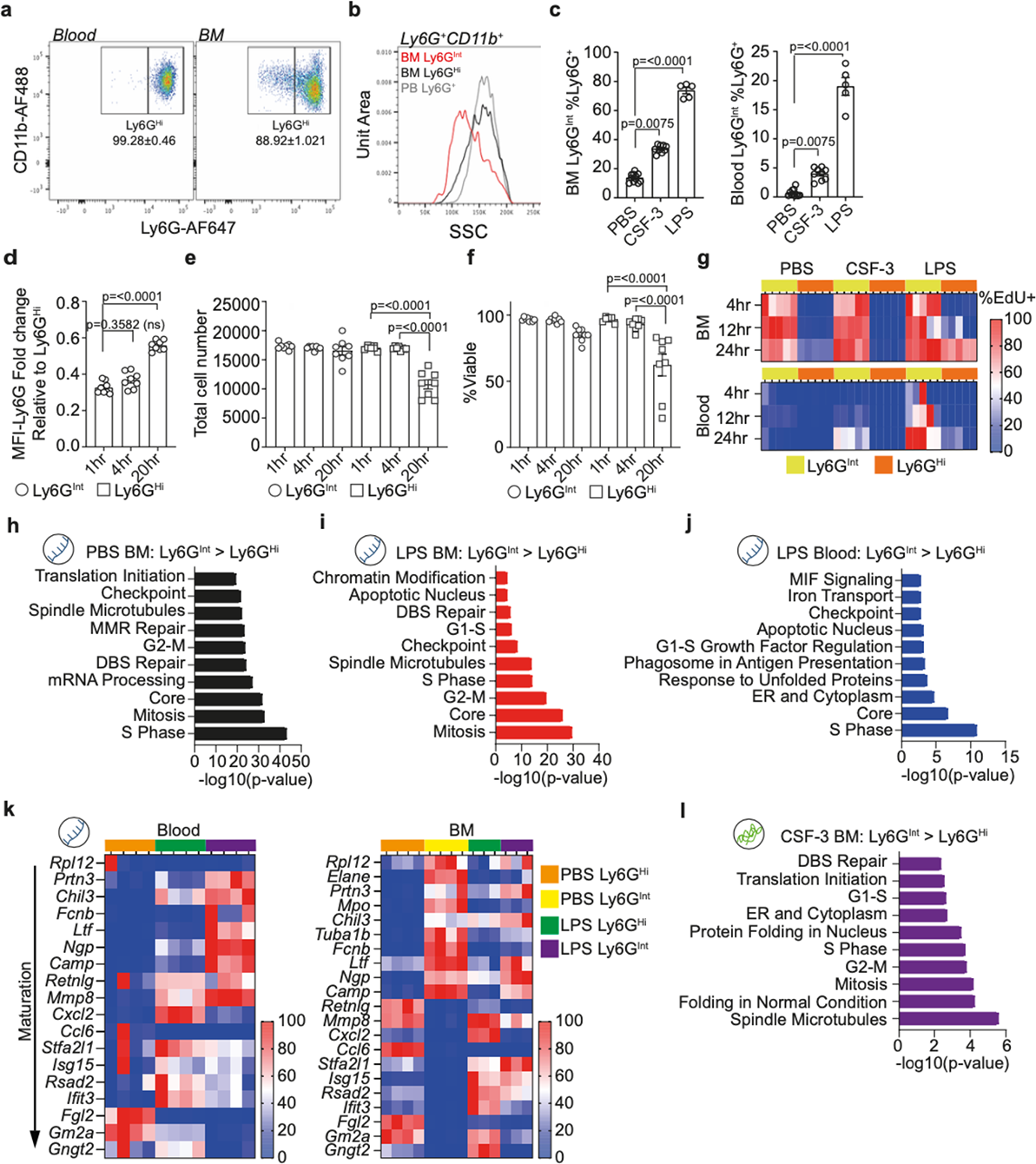
Intermediate surface Ly6G expression accurately and consistently identifies neutrophil immaturity. **a**, Flow cytometry plot for Ly6G^Int^ and Ly6G^Hi^ neutrophils in the peripheral blood and BM of naive mice. Data are presented as mean±standard deviation. Data representative of n=5 mice. **b**, Histogram showing flow cytometric SSC profile for BM Ly6G^Int^, BM Ly6G^Hi^ and peripheral blood Ly6G^+^ neutrophils from naïve mice. Data representative of n=5 mice. **c**, Quantification of Ly6G^Int^ as %Ly6G^+^ in the BM (left) and peripheral blood (right). PBS n=11 mice; CSF-3 n=9 mice; LPS n=5 mice. Data were analysed by Kruskal-Wallis test with Dunn’s multiple comparisons test. **d**, Quantification of MFI-Ly6G fold change for Ly6G^Int^ relative to Ly6G^Hi^ neutrophils isolated from naïve mice following *ex vivo* culture. 1hr time-point n=8 mice; 4hr time-point n=8 mice; 20hr time-point n=8 mice. Data were analysed by Kruskal-Wallis test with Dunn’s multiple comparisons test. **e**, Quantification of total cell no. recovered for Ly6G^Int^ and Ly6G^Hi^ neutrophils isolated from naïve mice following *ex vivo* culture. 1hr time-point n=8 mice; 4hr time-point n=8 mice; 20hr time-point n=8 mice. Data were analysed by 2way ANOVA with Tukey’s multiple comparisons test. **f**, Quantification of %Viable Ly6G^Int^ and Ly6G^Hi^ neutrophils isolated from naïve mice following *ex vivo* culture. 1hr time-point n=8 mice; 4hr time-point n=8 mice; 20hr time-point n=8 mice. Data were analysed by 2way ANOVA with Tukey’s multiple comparisons test. **g**, Heatmap showing row-scaled EdU^+^ as %Ly6G^Int^ and Ly6G^Hi^ neutrophils at 4, 12 and 24 hours post-EdU injection. BM: PBS 4hr, 12hr and 24hr Ly6G^Int^ and Ly6G^Hi^ n=5 mice; CSF-3 4hr, 12hr and 24hr Ly6G^Int^ and Ly6G^Hi^ n=5 mice; LPS 4hr, 12hr and 24hr Ly6G^Int^ and Ly6G^Hi^ n=5 mice. Peripheral blood: PBS 4hr, 12hr and 24hr Ly6G^Int^ and Ly6G^Hi^ n=5 mice; CSF-3 4hr, 12hr and 24hr Ly6G^Int^ and Ly6G^Hi^ n=5 mice; LPS 4hr, 12hr and 24hr Ly6G^Int^ and Ly6G^Hi^ n=5 mice. **h**, Top 10 most enriched process networks from DEGs with increased expression in BM Ly6G^Int^ compared to BM Ly6G^Hi^ neutrophils in PBS-control mice. Data from transcriptomic analysis. **i**, Top 10 most enriched process networks from DEGs with increased expression in BM Ly6G^Int^ compared to BM Ly6G^Hi^ neutrophils from LPS-challenged mice. Data from transcriptomic analysis. **j**, Top 10 most enriched process networks from DEGs with increased expression in peripheral blood Ly6G^Int^ compared to peripheral blood Ly6G^Hi^ neutrophils from LPS-challenged mice. Data from transcriptomic analysis. **k**, Heatmap showing row scaled expression for genes associated with neutrophil maturation (identified by Xie *et al.,* 2020) for Ly6G^Int^ and Ly6G^Hi^ neutrophils in the peripheral blood (left) and BM (right). Data from transcriptomic analysis. **l**, Top 10 most enriched process networks from proteins with increased abundance in BM Ly6G^Int^ compared to BM Ly6G^Hi^ neutrophils from CSF-3-challenged mice. Data from proteomic analysis. Bulk Ly6G^Int^ and Ly6G^Hi^ neutrophil RNA-Seq data in Fig. 2h-k. PBS BM: Ly6G^Int^ n=4 mice; Ly6G^Hi^ n=4 mice. LPS-BM: Ly6G^Int^ n=3 mice; Ly6G^Hi^ n=3 mice. PBS peripheral blood: Ly6G^Hi^ n=4 mice. LPS peripheral blood: Ly6G^Int^ n=4 mice; Ly6G^Hi^ n=4 mice. Bulk Ly6G^Int^ and Ly6G^Hi^ neutrophil proteomic data in Fig. 2l. CSF-3 BM Ly6G^Int^ n=3, and peripheral blood and BM Ly6G^Hi^ n=3 replicates. Each replicate composed of FACS isolated Ly6G^Int^ and Ly6G^Hi^ neutrophils pooled from n=4 mice. Dots in Fig. 2c-f represent individual mice. Error bars in Fig. 2c-f represent mean±SEM.

Differentially Expressed Genes (DEGs) increased in Ly6G^Int^ compared to Ly6G^Hi^ neutrophils were consistently enriched for cell cycle regulatory process networks in the BM and peripheral blood of PBS-control and LPS-challenged mice, confirming that these remain a more phenotypically immature population in the circulation (**Fig. 2h-j**). Using two recently identified gene signatures for stages of neutrophil development^8, 16^ we positioned Ly6G^Int^ neutrophils between the promyelocyte and metamyelocyte stages in PBS-control and LPS-challenged mice (**Fig. 2k** **and Extended Data Fig. 2g**). Corroborating this, BM Ly6G^Int^ neutrophils from CSF-3- challenged mice were enriched in proteins involved in cell cycle regulation (**Fig. 2l**). Thus, Ly6G surface expression accurately identifies immature Ly6G^Int^ and mature Ly6G^Hi^ neutrophils in homeostasis and models of haematopoietic stress.

### Other proposed markers of neutrophil maturity are altered in haematopoietic stress

Several other strategies have been proposed for measuring neutrophil maturity during homeostasis and/or particular disease models. We next sought to determine how generally applicable these markers are in non-tumour models by using CSF-3^22, 23^ and LPS-challenge^24^. Using Ly6G surface expression as a reference point for neutrophil maturity, we investigated whether Ly6G^Int^ neutrophils could be identified as tdTomato^Dim/-^ and Ly6G^Hi^ as tdTomato^Bright/+^ using Catchup^IVM-red^ mice in which *Ly6G-Cre* is used to drive tdTomato expression in neutrophils^25^. Ly6G^+^tdTomato^Bright/+^ cells composed a significantly higher percentage of Ly6G^Hi^ neutrophils compared to their Ly6G^Int^ counterparts in the peripheral blood and BM of all models (**Fig. 3a** **and Extended Data Fig. 3a**). However, CSF-3- and LPS-challenge significantly reduced the proportion of tdTomato^Bright/+^ neutrophils (**Extended Data Fig. 3a, b**). Here, Ly6G^Hi^ neutrophils were as low as 60% tdTomato^Bright/+^ in the peripheral blood of LPS-challenged mice (**Fig. 3a**). Consequently, tdTomato intensity was not specific enough to accurately determine neutrophil maturity during haematopoietic stress.

**Fig. 3.**
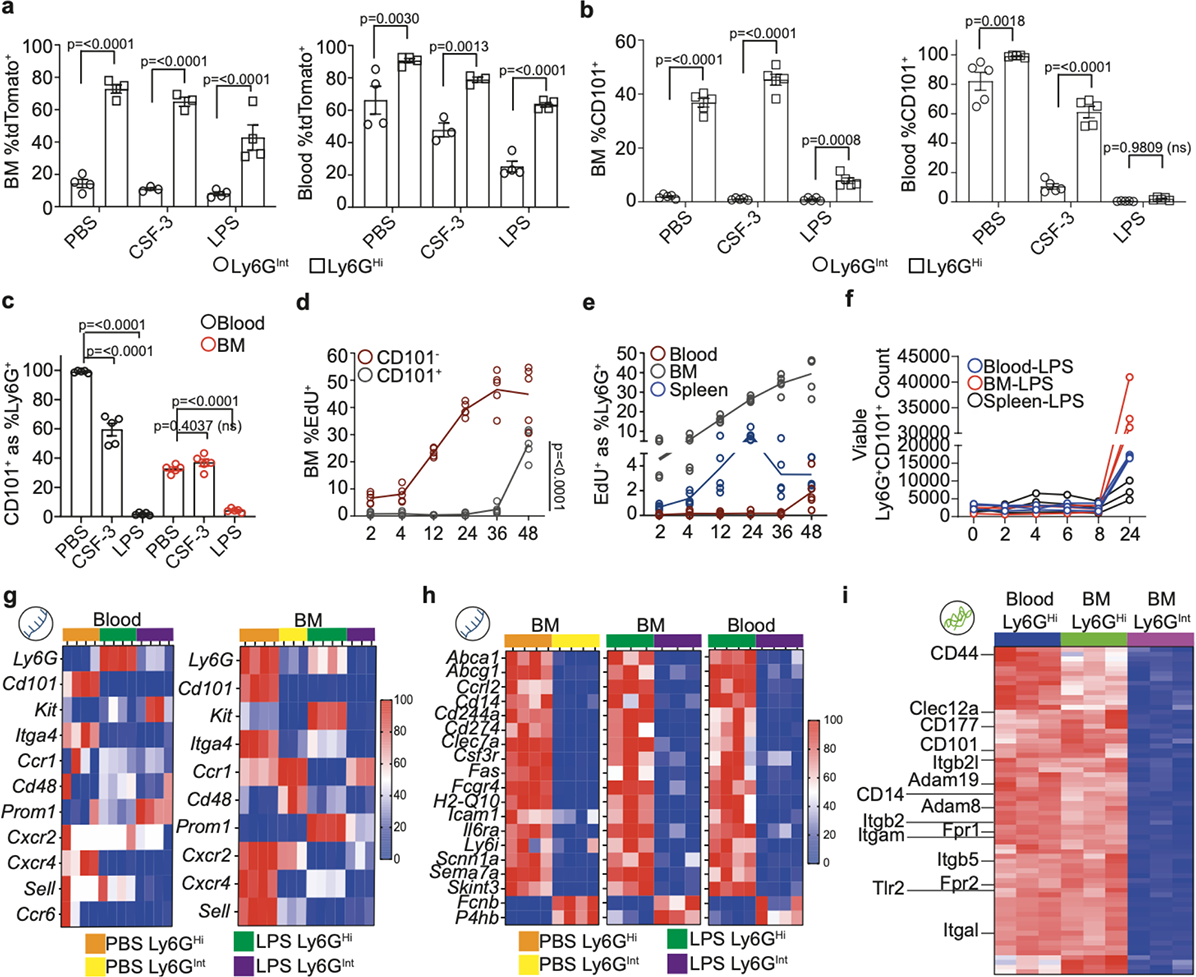
Alternative methods for identifying neutrophil maturity are less accurate in states of haematopoietic stress. **a**, Quantification of TdTomato^+^ as %Ly6G^Int^ and Ly6G^Hi^ neutrophils in the BM (left) and peripheral blood (right) of Catchup^IVM-red^ mice (developed by Hasenberg *et al.,* 2015). BM Ly6G^Int^ and Ly6G^Hi^ neutrophils: PBS n=4 mice; CSF-3 n=3 mice; LPS n=4 mice. Peripheral blood Ly6G^Int^ and Ly6G^Hi^ neutrophils: PBS n=4 mice; CSF-3 n=3 mice; LPS n=4 mice. Data from BM and peripheral blood were analysed by 2way ANOVA with Sidak’s multiple comparisons test. **b**, Quantification of CD101^+^ as %Ly6G^Int^ and Ly6G^Hi^ neutrophils in the BM (left) and peripheral blood (right). BM Ly6G^Int^ and Ly6G^Hi^ neutrophils n=5 mice in PBS, CSF-3 and LPS. Peripheral blood Ly6G^Int^ and Ly6G^Hi^ neutrophils n=5 mice in PBS, CSF-3 and LPS. Data from BM and peripheral blood were analysed by 2way ANOVA with Sidak’s multiple comparisons test. **c**, Quantification of CD101^+^ as %Ly6G^+^ neutrophils. Peripheral blood and BM n=5 mice in PBS, CSF-3 and LPS. Data were analysed by 2way ANOVA with Sidak’s multiple comparisons test. **d**, Quantification of EdU^+^ as %CD101^+^ in the BM of naïve mice at 2, 4, 12, 24, 36 and 48 hours post-EdU injection. CD101^-^ and CD101^+^ n=5 mice at all time-points. Data were analysed by 2way ANOVA with Tukey’s multiple comparisons test. **e**, Quantification of EdU^+^ as %Ly6G^+^ in the peripheral blood, BM and spleen of naïve mice at 2, 4, 12, 24, 36 and 48 hours post-EdU injection. Peripheral blood, BM and spleen Ly6G^+^ neutrophils n=5 mice at all time-points. **f**, Quantification of the no. of viable Ly6G^+^CD101^+^ neutrophils isolated from the peripheral blood, BM and spleen of LPS-challenged mice after 0, 2, 4, 6, 8 and 24 hours culture *ex vivo.* Peripheral blood, BM and spleen Ly6G^+^CD101^+^ neutrophils n=3 mice at all time-points. **g**, Heatmap showing row scaled expression for genes associated with proposed cell surface markers of neutrophil maturation. Data from transcriptomic analysis. **h**, Heatmap showing row scaled expression for genes with proteins localized on the external side of the plasma membrane (GO:0009897). Data from transcriptomic analysis. **i**, Heatmap showing row scaled protein abundance for genes whose proteins are localized within the cell membrane and are increased in Ly6G^Hi^ compared to Ly6G^Int^ neutrophils. Data from proteomic analysis. Bulk Ly6G^Int^ and Ly6G^Hi^ neutrophil RNA-Seq data in Fig. 3g, h. PBS BM: Ly6G^Int^ n=4 mice; Ly6G^Hi^ n=4 mice. LPS-BM: Ly6G^Int^ n=3 mice; Ly6G^Hi^ n=3 mice. PBS peripheral blood: Ly6G^Hi^ n=4 mice. LPS peripheral blood: Ly6G^Int^ n=4 mice; Ly6G^Hi^ n=4 mice. Bulk Ly6G^Int^ and Ly6G^Hi^ neutrophil proteomic data in Fig. 2l. CSF-3 BM Ly6G^Int^ n=3, and peripheral blood and BM Ly6G^Hi^ n=3 replicates. Each replicate composed of FACS isolated Ly6G^Int^ and Ly6G^Hi^ neutrophils pooled from n=4 mice. Dots in Fig. 3a-f represent individual mice. Error bars in Fig. 3a-c represent mean±SEM.

Cell surface CD101 expression has been proposed to distinguish between CD101^-^ immature and CD101^+^ mature neutrophils^10^. Consistent with this, a significantly greater percentage of Ly6G^Hi^ neutrophils were CD101^+^ compared to Ly6G^Int^ neutrophils in PBS-control mice (**Fig.3b and Extended Data Fig. 3c**). However, neutrophil surface CD101 expression was drastically reduced in CSF-3- and LPS-challenge (**Fig. 3b, c** **and Extended Data Fig. 3c**). EdU pulse-chase experiments identified BM neutrophil CD101 surface expression occurs 36-48 hours after the last cell division, the same time as the appearance of EdU^+^Ly6G^+^ in the peripheral blood (**Fig. 3d, e**). Thus, CD101 surface expression occurs during the terminal stages of neutrophil maturation and neutrophils can enter the blood prior to surface CD101 expression during haematopoietic stress. Despite this, when cultured *ex vivo,* CD101^-^ neutrophils from LPS-challenged mice upregulated CD101 surface expression between 8 and 24 hours, indicating that these neutrophils maintain the potential to express CD101 outside of this inflammatory context (**Fig. 3f**). Combined Ly6G and c-Kit (CD117) staining has also been used to identify neutrophil populations at distinct maturation stages in mice^14, 26^ (**Extended Data Fig. 3d**). CD117 surface expression is restricted to BM Ly6G^Int^ neutrophils in PBS-controls but is down-regulated in the BM and up-regulated by peripheral blood Ly6G^Hi^ neutrophils in CSF-3- and LPS- challenged mice (**Extended Data Fig. 3d**). Using RNA-Seq analysis we confirmed transcriptional regulation consistent with all of the findings above (**Fig. 3g**). Additionally, we identified stimulus dependent, variable gene and protein expression for other potential markers of neutrophil maturity (e.g. *Cd48*^10^*, Cd133*^27^) by Ly6G^Int^ and Ly6G^Hi^ neutrophils (**Fig. 3g** **and Extended Data Fig. 3e**).

We next attempted to identify novel candidate cell surface markers of neutrophil maturity. Using RNA sequencing (RNA-Seq) and gene ontology (GO) analysis, we found 19 DEGs between BM Ly6G^Int^ and Ly6G^Hi^ neutrophils in PBS-control and LPS-challenged mice whose proteins localise to the external side of the plasma membrane (GO:009897; **Fig. 3h**). Interrogating our proteomic dataset, we also identified 68 proteins that localized within the plasma membrane with different abundance between Ly6G^Int^ and Ly6G^Hi^ populations in CSF-3-challenged mice (**Fig. 3i**). However, the proteins identified included those that we and others have shown to be unreliable markers of neutrophil maturity dependent on stimuli, including CD177^28^, CXCR2^10^ and CD101. Therefore, as Ly6G expression closely matched objective measures of neutrophil maturity and remained our most reliable indicator compared to all others tested, we subsequently refer to Ly6G^Int^ neutrophils as ‘immature’ and Ly6G^Hi^ as ‘mature’ in the text.

### Immature neutrophils have reduced immune regulatory and adhesion capacity

As neutrophils develop effector mechanisms during maturation, for example neutrophil granules^29^, neutrophils at different maturation stages should have different functional capacity. Like transcriptomic differences (**Extended Data Fig. 2e, f**), proteomic analysis identified a striking difference in protein content of BM immature compared to BM and peripheral blood mature neutrophil populations (**Extended Data Fig. 4a**). Here, proteins increased in immature neutrophils were predominantly involved in DNA, RNA, metabolism/catabolism and biosynthesis/building associated biological processes, consistent with our RNA-Seq data (**Fig. 4a**). The proteins increased in mature neutrophils were preferentially involved in adhesion/motility, regulation/signalling and cellular response associated biological processes and process networks (**Fig. 4a, b**). LPS-challenge induced a clear shift in gene transcription by both immature and mature neutrophils compared to PBS-controls, including down-regulation (e.g. *Cd101, Ptgs2, Cxcr1*) and up-regulation (e.g. *Nos2, Ngp, Il10*) of genes (**Table 1,** **Fig. 4c** **and Extended Data Fig. 4b**). Despite this, the DEGs increased in LPS-challenge peripheral blood mature neutrophils were also preferentially involved in immune response regulation including phagocytosis and adhesion (**Extended Data Fig. 4c**).

**Fig. 4.**
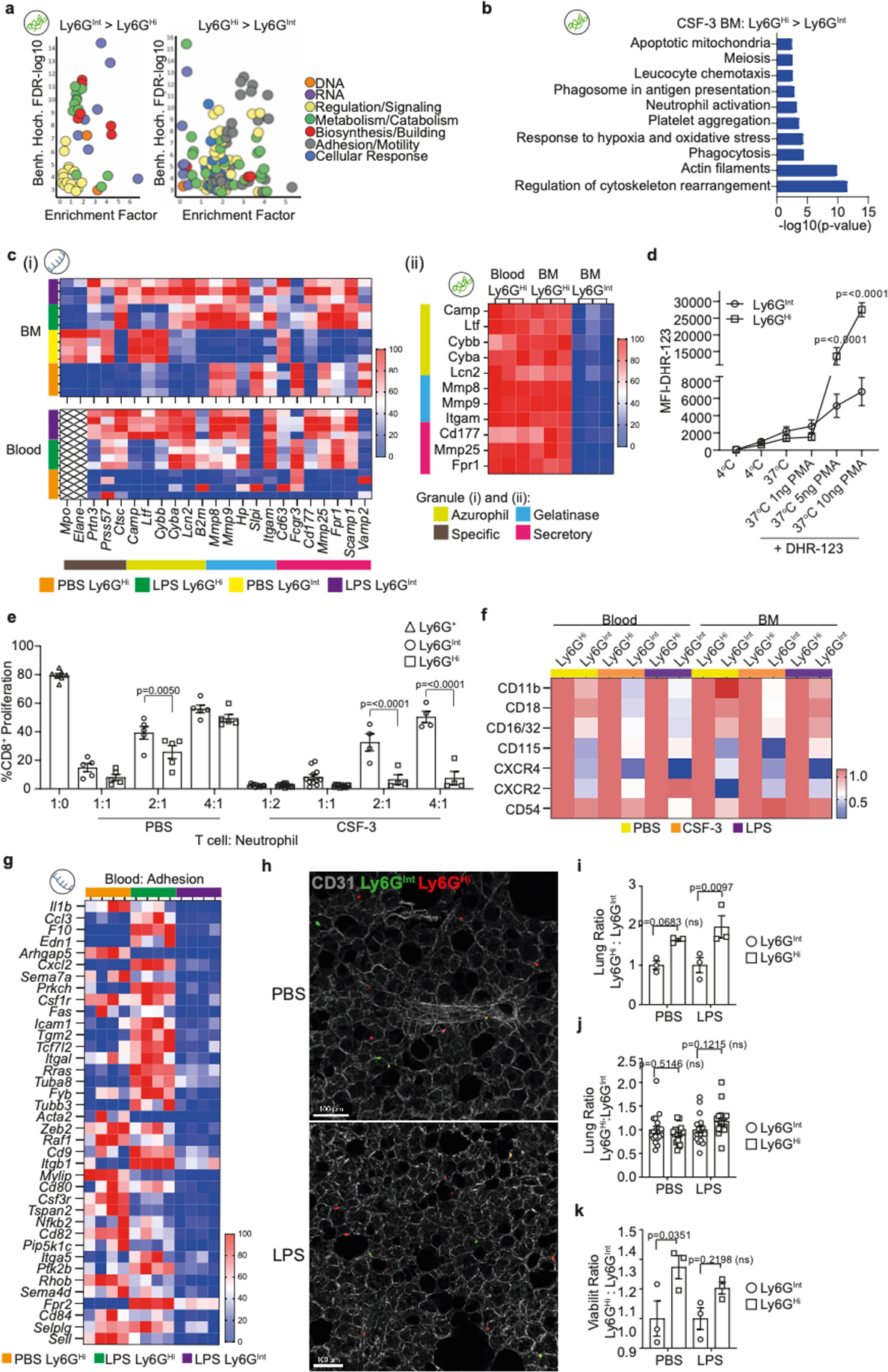
Ly6G^Int^ neutrophils have reduced functional capacity compared to Ly6G^Hi^. **a**, GO Biological Process clustering for proteins enriched in BM Ly6G^Int^ compared to Ly6G^Hi^ (left) and BM Ly6G^Hi^ compared to Ly6G^Int^ (right) from CSF-3-challenged mice. Data from proteomic analysis. **b**, Top 10 most enriched process networks from proteins with increased abundance in BM Ly6G^Hi^ compared to BM Ly6G^Int^ neutrophils from CSF-3-challenged mice. Data from proteomic analysis. **c**, Heatmap showing row-scaled (i) gene expression and (ii) protein abundance for neutrophil granule components. Data from (i) transcriptomic and (ii) proteomic analysis. **d**, Quantification of MFI for DHR-123 in *ex vivo* PMA stimulated Ly6G^Int^ and Ly6G^Hi^ neutrophils isolated from the BM of naïve mice. Data from flow cytometry analysis. Ly6G^Int^ and Ly6G^Hi^ neutrophils n=6 mice for each condition. Data were analysed by 2way ANOVA with Sidak’s multiple comparisons test. **e**, Quantification of %CD8^+^ T cell proliferation when cultured *ex vivo* with Ly6G^Int^ or Ly6G^Hi^ neutrophils isolated from the BM of PBS-control or CSF-3-challenged mice, at different T cell: neutrophil ratios. Data from flow cytometry analysis. Ly6G^+^ neutrophils 1:0 ratio n=7 mice. Ly6G^Int^ and Ly6G^Hi^ neutrophils: PBS 1:1, 2:1 and 4:1 ratios n=5 mice; CSF-3 1:2 and 1:1 ratios n=11 mice; CSF-3 2:1 and 4:1 ratios n=4 mice. Data were analysed by 2way ANOVA with Sidak’s multiple comparisons test. **f**, Heatmap showing row scaled CD11b^+^, CD18^+^, CD16/32^+^, CD115^+^, CXCR4^+^, CXCR2^+^ and CD54 as %Ly6G^Int^ and Ly6G^Hi^ neutrophils isolated from the peripheral blood and BM of PBS-control, CSF-3- and LPS-challenged mice. Data from flow cytometry analysis. Data are shown as ratio between Ly6G^Int^ and Ly6G^Hi^ where Ly6G^Hi^= 1. Peripheral blood and BM Ly6G^Int^ and Ly6G^Hi^ neutrophils: CD11b PBS n=8 mice, CSF-3 n=7 mice, LPS n=8 mice; CD18 PBS n= 5 mice, CSF-3 n=7 mice; LPS n=3 mice; CD16/32 PBS n=3 mice, CSF-3 n=3 mice, LPS n=3 mice; CD115 PBS n=3 mice, CSF-3 n=3 mice, LPS n=3 mice; CXCR4 PBS n=5 mice, CSF-3 n= 7 mice, LPS n=3 mice; CD54 PBS n=5 mice, CSF-3 n=4 mice, LPS n=5 mice; CXCR2 PBS n=3 mice, CSF-3 n=7 mice, LPS n=4 mice. **g**, Heatmap showing row-scaled expression for genes associated with leukocyte adhesion in peripheral blood Ly6G^Int^ and Ly6G^Hi^ neutrophils. Data from transcriptomic analysis. **h**, Confocal images of adoptively transferred Ly6G^Int^ and Ly6G^Hi^ neutrophils in the lungs of PBS-control and LPS-challenged recipient mice. Data representative of PBS n=3 mice and LPS n=3 mice. **i**, Quantification of adoptively transferred Ly6G^Int^ and Ly6G^Hi^ neutrophils recovered from the right lung lobe of recipient mice, expressed as a fold change per replicate mouse for Ly6G^Int^ over Ly6G^Hi^. Data from flow cytometry analysis. PBS n=3 mice; LPS n=3 mice. Data were analysed by 2way ANOVA with Sidak’s multiple comparisons test. **j**, Quantification of the number of adoptively transferred Ly6G^Int^ and Ly6G^Hi^ neutrophils identified in precision cut lung slices (PCLS) of recipient mice by confocal microscopy, expressed as a ratio between Ly6G^Int^: Ly6G^Hi^. PBS n=3 mice; LPS n=3 mice. From each mouse 2x images from 3x PCLS were analysed. Data were analysed by 2way ANOVA with Sidak’s multiple comparisons test. **k**, Viability of Ly6G^Int^ and Ly6G^Hi^ neutrophils recovered from the right lung lobe of recipient mice, expressed as a fold change per replicate mouse for Ly6G^Int^ over Ly6G^Hi^. Data from flow cytometry analysis. PBS n=3 mice; LPS n=3 mice. Data were analysed by 2way ANOVA with Sidak’s multiple comparisons test. Bulk Ly6G^Int^ and Ly6G^Hi^ neutrophil RNA-Seq data in Fig. 4c(i), g. PBS BM: Ly6G^Int^ n=4 mice; Ly6G^Hi^ n=4 mice. LPS-BM: Ly6G^Int^ n=3 mice; Ly6G^Hi^ n=3 mice. PBS peripheral blood: Ly6G^Hi^ n=4 mice. LPS peripheral blood: Ly6G^Int^ n=4 mice; Ly6G^Hi^ n=4 mice. Bulk Ly6G^Int^ and Ly6G^Hi^ neutrophil proteomic data in Fig. 2l. CSF-3 BM Ly6G^Int^ n=3, and peripheral blood and BM Ly6G^Hi^ n=3 replicates. Each replicate composed of FACS isolated Ly6G^Int^ and Ly6G^Hi^ neutrophils pooled from n=4 mice. Dots in Fig. 4d represent averages. Dots in Fig. 4e, i, k represent individual mice. Dots in Fig. 4j represent individual images from PCLS. Error bars in Fig. 4d, e, i-k represent mean±SEM.

Whereas BM immature neutrophils from PBS-control mice had increased transcription of azurophil and gelatinase granule-associated genes (**Fig. 4c(i)**), immature neutrophils from the BM and peripheral blood of LPS-challenged mice had increased transcription of genes associated with all granule types consistent with a more rapid maturation phenotype following LPS-challenge (**Fig. 4c(i)**). Consistently, proteomics showed that immature neutrophils had reduced levels of specific, gelatinase and secretory granule proteins (**Fig. 4c(ii)**).

We next sought to investigate the functional capacity of immature and mature neutrophils. Proteomic analysis revealed that NADPH-components were reduced in immature compared to mature neutrophils (**Extended Data** **Fig. 4d**). *Ex vivo* quantification of intracellular ROS with DHR-123 identified a slight increase in intracellular ROS production at 4°C, 37°C and 37°C with 1ng PMA stimulation by immature neutrophils (**Fig. 4d**). In contrast, the intracellular oxidative burst was far greater in mature neutrophils when stimulated with more PMA (**Fig. 4d** **and Extended Data Fig. 4e**). Neutrophils have been shown to suppress T cells in several disease contexts and maturity has been implicated in their suppressive capacity^4^. However, both immature and mature neutrophils from control mice suppressed CD8^+^ and CD4^+^ T cell proliferation and reduced their viability in a ratio-dependent manner (**Fig. 4e** **and Extended Data Fig. 4f-h**). This effect was stronger when using mature compared to immature neutrophils and was further augmented when T cells were cultured with mature neutrophils isolated from the BM of CSF-3-treated mice (**Fig. 4e** **and Extended Data Fig. 4g, h**). This suggests that the environment the cells are taken from plays a role in this behaviour rather than maturity alone, and that immature and mature neutrophils can have a different functional response to the same environment (**Fig. 4e** **and Extended Data Fig. 4g, h**).

We found that cell membrane localized adhesion and cell-cell interaction associated (e.g. ITGAM, CD11b; ITGAL, CD11a; ITGB2, CD18; ITGB2L; ITGB5; CD177; CD44; CLEC12a; ADAM19; ADAM8) and pathogen detection (e.g. CD14, TLR2, FPR1, FPR2) proteins were highly represented in mature neutrophils (**Fig. 3i** **and** **Fig. 4f**). In the peripheral blood of LPS-challenged mice, immature neutrophils had reduced gene expression of adhesion associated genes compared to mature ones (**Fig. 4g**). Flow cytometry confirmed lower surface expression of adhesion molecules (CD11b, CD18, CD54) and chemokine receptors (CXCR2, CXCR4) by immature neutrophils (**Fig. 4f**). These data are consistent with immature neutrophils having a reduced/altered effector capacity compared to their mature counterparts and emphasise their plasticity to develop the proteins needed during maturation, and also in response to stimuli, rather than being equipped with a fixed set at early stages. We next sought to test for consequent reduced adhesion of immature neutrophils *in vivo*. During sepsis, neutrophil retention in the lung can promote acute respiratory distress syndrome (ARDS), worsening pathology^30^. In mice, systemic LPS-challenge is commonly used to model sepsis-induced ARDS^31, 32^. To obtain large enough numbers, BM-derived immature and mature neutrophil populations were sorted from CSF-3-challenged mice and intravenously (i.v.) adoptively transferred into PBS-control or LPS-challenged mice. Fewer adoptively transferred immature neutrophils were detected in the lungs of PBS-control and LPS-challenged mice compared to mature neutrophils when analysed by flow cytometry and microscopy of precision cut lung slices (**Fig. 4h-j**). Furthermore, adoptively transferred immature neutrophils had reduced viability compared to mature neutrophils (**Fig. 4k**). Together, these data indicate that immature neutrophils are less capable at adhesion, migration, trafficking, pathogen removal, ROS production, and T cell suppression based on transcriptomic, proteomic, and flow cytometric, in addition to *in vivo* and *ex vivo* functional assays.

### Homeostatic neutrophil development in the spleen is distinct from the BM

Neutrophil tissue localization can influence their gene expression^3, 9^. Consistent with this, when we compared neutrophils isolated from separate tissues, immature and mature neutrophils had unique transcriptional profiles compared to the same maturation stage in other sites (**Table 2-4 and** **Fig. 5a**). Interestingly, while LPS-challenge induced a clear shift in the expression profile of both immature and mature neutrophils isolated from the same tissue compared to PBS-controls (PC2 variance 27.4%), variance between neutrophils isolated from separate tissues was greater (PC1 variance 49.5%; **Fig. 5a**). Furthermore, tissue variance was only slightly reduced in LPS-challenged mice compared to PBS-controls, indicating tissue specific signatures are largely maintained in acute inflammation (**Fig. 5a**).

**Fig. 5.**
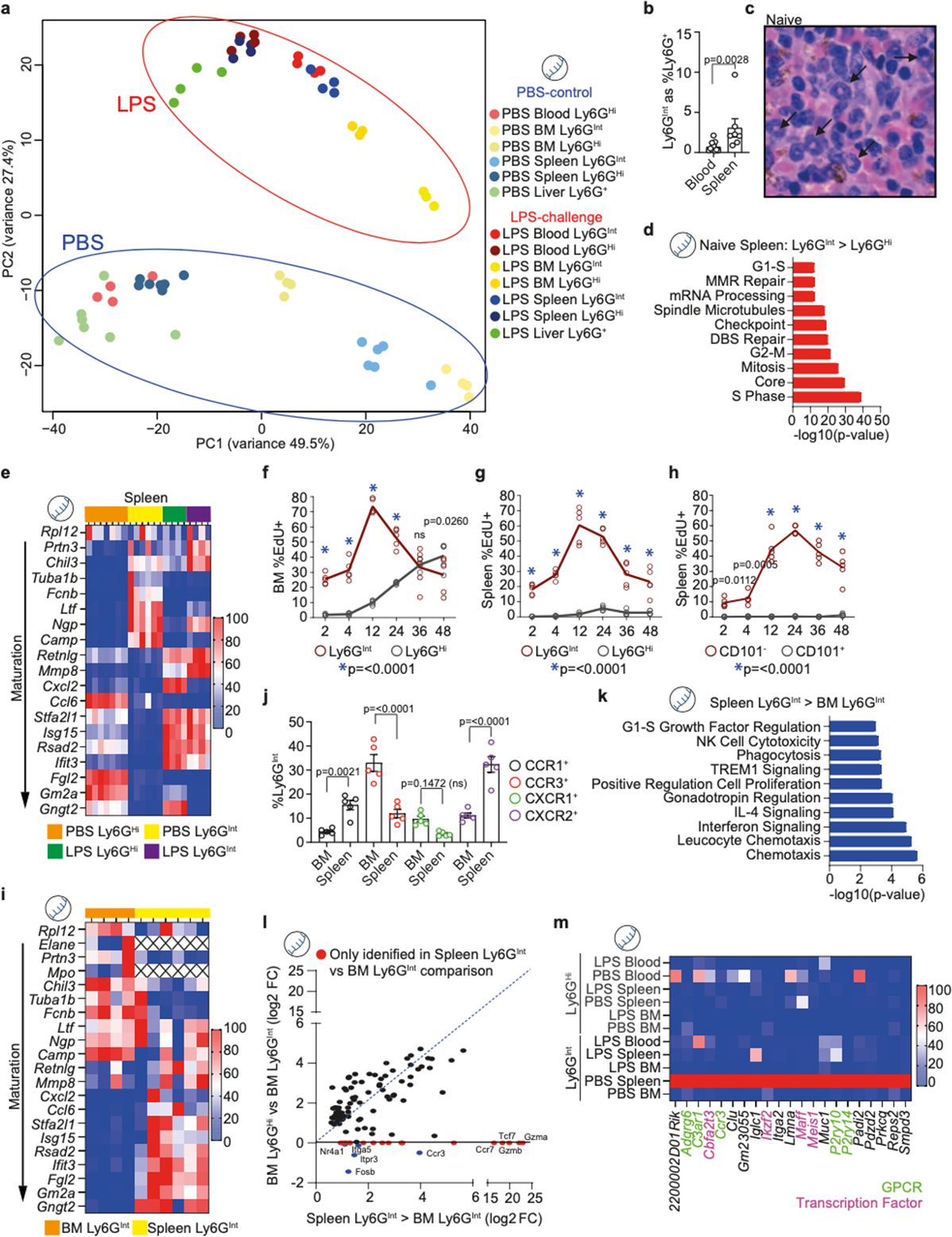
Splenic neutrophil maturation is distinct from the BM. **a**, PCA plot for Ly6G^+^, Ly6G^Int^ and Ly6G^Hi^ transcriptomic analysis from the peripheral blood, BM, spleen and liver of PBS-control and LPS-challenged mice. **b,** Quantification of Ly6G^Int^ as %Ly6G^+^ in the peripheral blood and spleen of naïve mice. Peripheral blood n=11 mice; spleen n=7 mice. Data were analysed by Mann-Whitney test. **c**, H&E identification of banded neutrophils in the splenic red pulp of naïve mice. Data representative of n=5 mice. Black arrows indicate banded neutrophils. **d**, Top 10 most enriched Process Networks from DEGs with increased expression in spleen Ly6G^Int^ compared to spleen Ly6G^Hi^ neutrophils from naive mice. Data from transcriptomic analysis. **e**, Heatmap showing row scaled expression for genes associated with neutrophil maturation (identified by Xie *et al.,* 2020) for Ly6G^Int^ and Ly6G^Hi^ neutrophils in spleen. Data from transcriptomic analysis. **f**, Quantification of EdU^+^ as %Ly6G^Int^ and Ly6G^Hi^ neutrophils in the BM of naïve mice. Ly6G^Int^ and Ly6G^Hi^ neutrophils n=5 mice at each time-point. Data were analysed by 2way ANOVA with Sidak’s multiple comparisons test. Blue star (*) indicates comparisons with p-value<0.0001. For ‘ns’ at 36hr time-point, p-value=0.9989. **g**, Quantification of EdU^+^ as %Ly6G^Int^ and Ly6G^Hi^ neutrophils in the spleen of naïve mice. Ly6G^Int^ and Ly6G^Hi^ neutrophils n=5 mice at each time-point. Data were analysed by 2way ANOVA with Sidak’s multiple comparisons test. Blue star (*) indicates comparisons with p-value<0.0001. **h**, Quantification of EdU^+^ as %CD101^-^ and CD101^+^ neutrophils in the spleen of naïve mice. CD101^-^ and CD101^+^ neutrophils n=5 mice at each time-point. Data were analysed by 2way ANOVA with Sidak’s multiple comparisons test. Blue star (*) indicates comparisons with p-value<0.0001. **i**, Heatmap showing row scaled expression for genes associated with neutrophil maturation (identified by Xie *et al.,* 2020) for Ly6G^Int^ neutrophils in BM and spleen from naïve mice. Data from transcriptomic analysis. **j**, Quantification of CCR1^+^, CCR3^+^, CXCR1^+^ and CXCR2^+^ as %Ly6G^Int^ neutrophils in the BM and spleen of naïve mice. Data from flow cytometry analysis. BM Ly6G^Int^ n=5 mice; spleen Ly6G^Int^ n=5 mice. Data were analysed by 2way ANOVA with Sidak’s multiple comparisons test. **k**, Top 10 most enriched Process Networks from DEGs with increased expression in spleen Ly6G^Int^ compared to BM Ly6G^Int^ neutrophils from naive mice. Data from transcriptomic analysis. **l**, Correlation for genes identified in process networks from Fig. 5k between spleen Ly6G^Int^ and BM Ly6G^Hi^, shown as log2 fold change (FC) compared to BM Ly6G^Int^. Data from transcriptomic analysis. **m**, Heatmap showing column scaled expression for genes with high expression unique to spleen Ly6G^Int^ neutrophils. Data from transcriptomic analysis. Bulk Ly6G^+^, Ly6G^Int^ and Ly6G^Hi^ neutrophil RNA-Seq data in Fig. 5a, d, e, i, k-m. PBS: peripheral blood Ly6G^Hi^ n=4 mice; BM Ly6G^Int^ n=4 mice; BM Ly6G^Hi^ n=4 mice; spleen Ly6G^Int^ n=6 mice; spleen Ly6G^Hi^ n=7 mice; liver Ly6G^+^ n=7 mice. LPS: peripheral blood Ly6G^Int^ n=4 mice; peripheral blood Ly6G^Hi^ n=4 mice; BM Ly6G^Int^ n=3 mice; BM Ly6G^Hi^ n=3 mice; spleen Ly6G^Int^ n=4 mice; spleen Ly6G^Hi^ n=4 mice; liver Ly6G^+^ n=4 mice. Dots in Fig. 5a, b, f-h, j represent individual mice. Error bars in Fig. 5b, f-h, j represent mean±SEM.

Immature neutrophils were enriched in the spleen compared to other peripheral tissues and peripheral blood in control animals (**Fig. 5b**, **Fig. 1l-p** **(WT mice) and Extended Data Fig. 5a**) consistent with other reports of their presence in a steady state^5^. In the spleen of naïve mice, we found clusters of morphologically banded neutrophils localized in the red-pulp and using flow cytometry we identified unipotent neutrophil progenitors, termed NePs^33^ (**Fig. 5c** **and Extended Data Fig. 5b**). We next investigated whether tissue specific factors altered splenic neutrophil maturation compared to the BM. Similarly to the BM, splenic immature neutrophils had a different expression profile to mature neutrophils, enriched DEGs preferentially involved in cell cycle regulation compared to mature neutrophils, and were positioned between myelocyte and metamyelocytes using the 24 maturation stage-associated genes as above^16^ (**Fig. 5d, e** **and Extended Data Fig. 5c**). Additionally, as we had seen before in the BM, LPS-challenge increased expression of genes associated with later maturation stages, and the immature splenic neutrophils rapidly incorporated EdU within 2-4 hours of its injection, and contained Ki67^+^ cells in PBS-control, CSF-3- and LPS- challenged mice (**Fig. 5e-g** **and Extended Data Fig. 5d, e**).

In the BM, EdU^+^ immature neutrophils increased from 2-12 hours and reduced from 12-48 hours post-EdU pulse, coincident with an increase in EdU^+^ mature neutrophils consistent with Ly6G surface upregulation with maturation (**Fig. 5f**). However, although splenic EdU^+^ immature neutrophils increased from 2-12 hours and reduced between 12-48 hours post-injection, there was no coincident increase in EdU^+^ mature neutrophils (**Fig. 5g**). As in the BM CD101 surface expression occurred during the terminal stages of neutrophil maturation (**Fig. 3d**) we analysed CD101 expression in the spleen to explore whether splenic post-mitotic transit time was shorter when compared to the BM. We observed no significant increase in EdU^+^CD101^+^ neutrophils in the spleen up to 48 hours post-injection and only observed EdU^+^CD101^+^ neutrophils in the peripheral blood between 36-48 hours corresponding with their emergence from the BM (**Fig. 5h** **and Extended Data Fig. 5f**). As the spleen also clears neutrophils from the circulation^34^, we analysed pro-apoptotic genes (GO:0043065) in splenic immature neutrophils compared to other populations. As expected, mature neutrophils generally expressed these genes at higher levels than immature, but specifically, immature neutrophils in the spleen were not enriched for these suggesting that they do not apoptose there (**Extended Data** **Fig. 5g**). The BM contains a neutrophil reserve composed of mature, terminally differentiated neutrophils that can be rapidly mobilized in response to stress. The spleen has been proposed to contain marginated neutrophils, however, we found no evidence in the literature for a distinct splenic-neutrophil reserve analogous to that in the BM^35^. Therefore, we interpret these data as confirming the presence of an abundant splenic immature neutrophil population that is released earlier from its developmental niche, compared to those in the BM, which enter the BM reserve.

### Splenic-derived neutrophils have quicker maturation and altered transcriptional phenotype

Recently developed mature neutrophils form the BM-reserve and undergo important changes associated with their functional capacity, including their development of secretory vesicles^7^. To investigate whether maturation was faster in splenic-compared to BM-derived neutrophils to account for the absence of a splenic-reserve, we compared BM and spleen immature neutrophil gene expression profiles (**Table 2**). Splenic immature neutrophils had increased expression of later stage maturation genes compared to BM immature neutrophils in PBS-control mice (**Fig. 5i** **and Extended Data Fig. 5h**). Furthermore, spleen immature neutrophils had significantly different mRNA levels for several chemokine receptors (**Extended Data Fig. 5i**). Flow cytometry confirmed a significant increase in CCR1 and CXCR2, and a significant decrease in CCR3 surface expression, by splenic immature compared to BM immature neutrophils in naïve mice (**Fig. 5j**). In mature neutrophils, the percentage CCR1^+^ and CXCR2^+^ cells increased, and CCR3^+^ decreased, from BM to spleen to peripheral blood respectively (**Extended Data Fig. 5j**). When we directly compared BM and spleen immature neutrophils, we found those from BM were enriched in genes associated with cell cycle regulation and those from the spleen were enriched in inflammatory/immune response networks (**Fig. 5k** **and Extended Data Fig. 5k**). This closely matched genes associated with these process networks that were enriched in BM mature neutrophils compared to BM immature neutrophils, including interferon signalling (**Fig. 5l** **and Extended Data Fig. 5l**). However, expression of certain genes (*Nr4a1, Itga5, Itpr3, Ccr3, Fosb*) increased in spleen immature neutrophils had reduced expression in the BM mature-immature comparison and some were unique to spleen immature neutrophils (e.g. *Gzma, Gzmb, Tcf7, Ccr7*; **Fig. 5l**). We also identified several genes with high expression unique to PBS-control spleen immature neutrophils including GPCRs (e.g*. C3ar1, Pr2y10*) and transcription factors (e.g. *Ikzf2, Maff*; **Fig. 5m**). Together, these data are consistent with a quicker maturation phenotype, shorter post-mitotic transit and earlier release from the splenic developmental niche compared to neutrophils that originate in the BM, in addition to a unique splenic developmental-niche driven expression signature.

### Splenic-granulopoiesis remains distinct from the BM in tumour bearing mice

As previously shown, immature neutrophils were identified in the spleen of DEN/ALIOS, KPN, KC, KP^fl^C and KPC mice (**Fig. 1**). This was confirmed histologically by clusters of banded neutrophils localized in the splenic red-pulp (**Fig. 6a**). As splenic immature neutrophils were abundant in both highly metastatic models analysed, we next investigated if the presence of a PDAC and metastatic potential influences BM and spleen immature neutrophil phenotype. RNA-Seq of BM and splenic neutrophils isolated from WT, KP^fl^C and KPC mice confirmed an enrichment in genes associated with cell cycle regulation in BM and spleen immature compared to mature neutrophils (**Fig. 6b** **and Extended Data Fig. 6a**). The presence of a PDAC led to a dramatic shift in expression in BM and spleen immature neutrophils associated with an increased activated degranulatory phenotype (**Table 5, 6;** **Fig. 6c**; **Extended Data Fig. 6b**). However, direct comparison of BM and spleen immature neutrophils between KP^fl^C and KPC genotypes identified 1 (*Eps8l1*) and 0 DEGs, respectively (**Table 5, 6**). Despite this, in the BM we identified 88 DEGs unique to KP^fl^C, 267 DEGs unique to KPC and 187 DEGs shared by KP^fl^C and KPC BM immature neutrophils compared to those in WT mice (**Table 7**). In the spleen we identified 204 DEGs unique to KP^fl^C, 916 DEGs unique to KPC and 504 DEGs shared by KP^fl^C and KPC spleen immature neutrophils compared to WT (**Table 7**). We interpret the absence of DEGs between KP^fl^C and KPC, but increased number of DEGs between KPC compared to WT, as indicating a progressive gene expression profile from KP^fl^C to KPC in both the BM and the spleen (**Extended Data Fig. 6c**). Immature neutrophils from the BM and spleen of KPC mice have increased neutrophil activation associated gene expression over KP^fl^C compared to WT mice (**Fig. 6d**). Thus, while metastatic potential enhances splenic granulopoiesis it does not significantly alter the splenic or BM developmental phenotype, rather the presence of a PDAC promotes an enhanced neutrophil activation developmental phenotype.

**Fig. 6.**
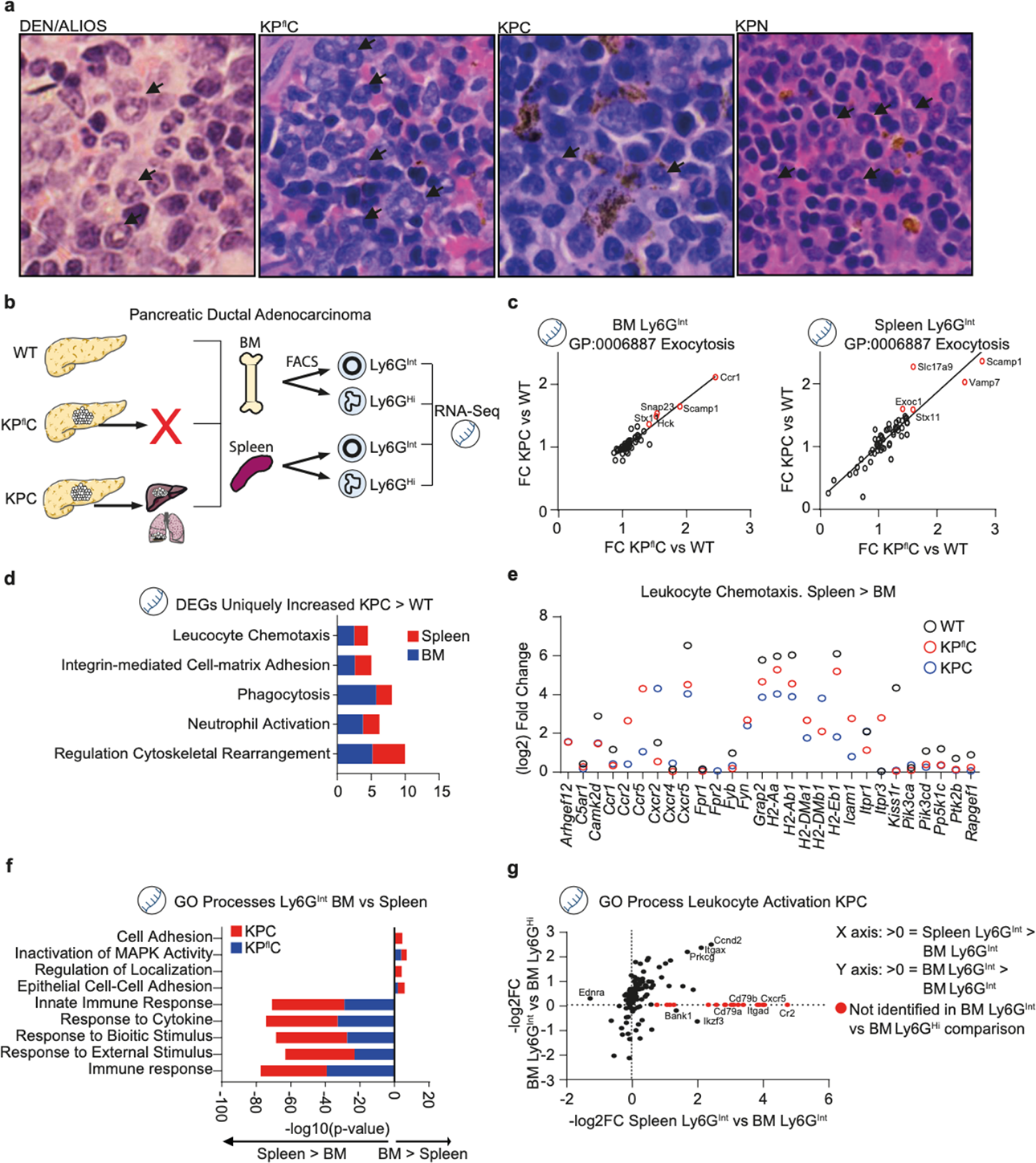
Splenic Ly6G^Int^ neutrophils maintain a distinct maturation phenotype in tumour bearing mice. **a**, H&E identification of banded neutrophils in the splenic red pulp of DEN/ALIOS, KP^fl^C, KPC and KPN mice. Data representative of DEN/ALIOS n=5 mice; KP^fl^C n=5 mice, KPC n=5 mice and KPN n=5 mice. **b**, Schematic for RNA-Seq of BM and spleen Ly6G^Int^ and Ly6G^Hi^ neutrophil populations from WT controls, KP^fl^C non-metastatic and KPC pre-metastatic mouse models of PDAC. **c**, Correlation for genes associated with exocytosis (GO:0006887) between KP^fl^C and KPC Ly6G^Int^ neutrophils in the BM (left) and spleen (right), shown as fold change compared to WT Ly6G^Int^ neutrophils. Data from transcriptomic analysis. **d**, Process Networks associated with neutrophil activation enriched from DEGs uniquely increased in KPC compared to WT BM and spleen. Data from transcriptomic analysis. **e**, DEGs associated with leukocyte chemotaxis with increased expression in spleen Ly6G^Int^ compared to BM Ly6G^Int^ in WT, KP^fl^C and KPC mice, shown as Log2 fold change. Data from transcriptomic analysis. **f**, GO Processes enriched from DEGs identified between KP^fl^C and KPC BM Ly6G^Int^ and spleen Ly6G^Int^ neutrophils. Data from transcriptomic analysis. **g**, Correlation for genes associated with leukocyte activation (GO:0045321) between spleen Ly6G^Int^ and BM Ly6G^Hi^, shown as Log2 fold change compared to BM Ly6G^Int^. Data from transcriptomic analysis. Bulk Ly6G^Int^ and Ly6G^Hi^ neutrophil RNA-Seq data in Fig. 5c-g. BM: WT Ly6G^Int^ n=10 mice; KP^fl^C Ly6G^Int^ n=5 mice; KPC Ly6G^Int^ n=5 mice; KPC Ly6G^Hi^ n=5 mice. Spleen: WT Ly6G^Int^ n=8 mice; KP^fl^C Ly6G^Int^ n=4 mice; KPC Ly6G^Int^ n=5 mice.

We next examined if BM and spleen immature neutrophils maintain their steady state differences in PDAC bearing mice. Splenic immature neutrophils from WT, KP^fl^C and KPC mice had an increased expression of chemotaxis associated genes and advanced maturation signature compared to BM immature neutrophils (**Fig. 6d, e** **and Extended Data Fig. 6d**). However, the presence of a PDAC reduced the total number of DEGs between BM and spleen immature neutrophils in part due to reduced expression of developmental associated genes in BM immature neutrophils (**Table 8 and Extended Data Fig. 6e, f**). Despite this, spleen immature neutrophils maintained an increase in immune responsiveness gene expression compared to those in the BM of PDAC-bearing mice (**Fig. 6f**). Similar to steady state, BM mature, and spleen immature neutrophils, had a more similar expression profile to each other when compared to BM immature neutrophils, such as for those involved in leukocyte activation (e.g. *Ccand2, Itgax, Prkcg*), while many genes were unique to spleen immature neutrophils (e.g. *Cxcr5, Itgad*; **Fig. 6g**). Thus, as in homeostasis, spleen immature neutrophils maintain a more mature expression profile compared to BM immature neutrophils with some distinct characteristics compared to both BM immature and mature populations. We next investigated if BM and spleen immature neutrophils undergo different changes in PDAC bearing mice when compared to WT. DEGs shared by KP^fl^C and KPC compared to WT from each tissue included 65 shared, 122 and 438 DEGs unique to BM and spleen immature neutrophils, respectively (**Extended Data Fig. 6g**). Here, the presence of a PDAC promoted an enhanced activated expression signature of splenic immature compared to BM immature neutrophils including increased expression of genes associated with antigen presentation (*Cd74, Cd79*) and immunosuppression (*Cd274, Btla*; **Extended Data Fig. 6h**). Therefore, differences likely to be functionally important between BM and spleen immature neutrophils are maintained in cancer models, but importantly, the increased abundance of these cells in cancer may enhance their functional consequences.

## Discussion

Neutrophils are promising targets for therapeutic intervention^36, 37^, however, due to their importance in antimicrobial defence, it is vital to target these therapies only at the neutrophils that are detrimental in disease. Immature neutrophils have been implicated in disease progression in a range of pathologies including sepsis and cancer^11, 15^ where they have been associated with immune suppression^4^, tumour burden^10^ and metastasis^38^. Thus, characterising immature neutrophils and their contribution to neutrophil heterogeneity is vital to efficiently targeting neutrophils in disease. We found that neutrophils expressing intermediate levels of Ly6G, variably abundant in poorly metastatic mouse cancer models, were enriched in two different spontaneously metastatic models. In the present study, we show that Ly6G expression level accurately identifies neutrophil immaturity by morphological, flow cytometric, transcriptomic, and proteomic analysis. This identification maintained its accuracy in homeostasis, experimental acute haematopoietic stress and chronic haematopoietic stress caused by cancer, across multiple tissues, and between different mouse strains, in contrast to other proposed maturity markers. Immature neutrophils have been proposed to have altered function compared to their mature counterparts as a direct result of their developmental status, including metabolic^14, 38^, immuno-suppressive^39^ and migratory^10^ capacity. Immature (Ly6G^Int^) neutrophils had reduced granule composition, ROS production following activation, suppressive capacity, cytoskeletal remodelling, and adhesion- and recruitment-associated proteins compared to mature (Ly6G^Hi^) neutrophils. Accordingly, fewer immature neutrophils were retained in the lungs of PBS- and LPS-challenged mice confirming reduced adhesion with potential implications in disease progression.

Extramedullary haematopoiesis (EMH) occurs physiologically in mice and is observed pathologically in both mice and humans. Consistent with the literature^5^, spleen immature neutrophils had increased proliferative and reduced immune response capacity compared to spleen mature neutrophils. However, our data positioned spleen immature neutrophils at a similar maturation stage to mature neutrophils in the BM reserve. Furthermore, increased expression of chemokine receptors, release prior to CD101 surface expression, and reduced cell cycle-associated gene expression are consistent with faster neutrophil maturation in the spleen than in the BM. Tissue localization can influence the neutrophil transcriptome^3^ and neutrophils can display tissue specific properties in mice and humans, including in the spleen^40^. We found that when compared to those from the BM, spleen immature neutrophils had distinct gene signatures that were associated with altered immune and transcriptional regulation. Thus, the site of haematopoietic origin influences neutrophil transit, maturation and functional characteristics even in a steady state. We propose that these data signify that spleen immature neutrophils are pre-potentiated ‘rapid responders’ following insult, more akin to reserve mature than immature neutrophils in the BM. It is anticipated that circulation of BM-derived progenitors in a steady state^41^ and disease promotes their seeding in the spleen, enhancing extramedullary myelopoiesis^42^. Differences observed between BM- and spleen-derived neutrophils likely arose from niche-induced alterations to development, similar to tissue-specific neutrophil phenotypes^3, 43, 44^.

Spleen-derived neutrophils contribute to the systemic pool in sterile chronic inflammation^45^ and cancer where they can acquire an immunosuppressive state^44, 46^. The spleen is a major site of extramedullary myelopoiesis in certain cancers and offers a potential therapeutic target^47^. Splenic-derived neutrophils can promote tumour progression^44^ and splenectomy can reduce metastatic burden^48^. We found that neutrophils were markedly increased in the peripheral blood and spleen, occupying large clusters in the splenic red-pulp, of highly metastatic CRC and PDAC mouse models. Here, we found that splenic immature neutrophils maintained their distinct expression phenotype associated with increased chemotaxis and immune responsiveness compared to BM immature neutrophils. Therefore, as metastasis is closely associated with patient mortality in cancer^49^ we postulate an important role for these abundant immature and splenic-derived neutrophils in cancer progression. Our data also indicate that developmental origin is critical in controlling neutrophil maturation and determining phenotype and function. However, EMH can occur at multiple sites in cancer^50^ and in other pathologies^51^ raising the potential for a plethora of phenotypically and functionally distinct neutrophil populations, on which deeper information is critical for the more specific targeting of neutrophils in disease.

## Methods

### Mouse strains

All mice were housed in specific pathogen free conditions with unrestricted access to food and water and maintained on a constant 12hr light/dark cycle under controlled climate (19-22°C, 45-65% humidity) at the CRUK Beatson Institute, Glasgow or Imperial College London. All experiments involving mice were performed in accordance with the UK Animal (Scientific Procedures) Act 1986, approved by the local animal welfare (AWERB) committee and conducted under UK Home Office licences: 70/7463; 70/8377; PP3908577; PE47BC0BF; PP6345023, and were compliant with the ARRIVE guidelines (https://www.nc3rs.org.uk/arrive-guidelines). For all experiments using WT C57BL/6 mice, 8-12 week old mice were used. C57BL/6 mice were purchased from Charles River Laboratories. Catchup^IVM-red^ mice, as first described^25^, were identified as *Ly6G*^Cre-Rtom/+^ *ROSA26*^Rtom/+^ and were bred in-house and used for experiments when they were 8-15 weeks old. For the metastatic model of CRC, KPN (*Villin*Cre^ER^; *Kras*^G12D/+^; *Trp53*^fl/fl^; *Rosa26*^N1icd/+^) mice^17^, were used. Mice of both sexes were induced with a single dose of 2mg tamoxifen (Sigma-Aldrich) by i.p. injection at 6-12 weeks old. Mice were sampled at humane clinical endpoint by weight loss and/or hunching and/or cachexia. For the DEN/ALIOS model of HCC, male WT C57BL/6 mice received a single dose of diethylnitrosamine (DEN; 80mg/kg) by intraperitoneal (i.p.) injection at 14 days old and were subsequently fed an American Lifestyle Induced Obesity Syndrome (ALIOS) diet (Envigo, TD.110201) from day 46 post-Den. Tumours were allowed to develop before mice were humanely killed at day 270 post-Den. For the poorly-metastatic model of lung adenocarcinoma, 8-10 week old KM (*<u>K</u>Ras*^G12D/+ 52^*; Rosa26^lsl-<u>M</u>yc/lsl-Myc 18^* ) mice were induced with 1x10^7^ pfu Adeno-*Cre* recombinant adenovirus (University of Iowa gene therapy core facility), as described previously. Induced KM mice and un-induced litter mates were humanely killed after 6 weeks when mice displayed no clinical signs but had macroscopic lesions^18^, and tissues harvested for analysis. For highly-metastatic and poorly-metastatic models of PDAC and their controls, WT, *<u>K</u>ras*^G12D^; *Pdx1*^Cre^ (KC), *<u>K</u>ras*^G12D^; *Trp53*^fl/+^; *Pdx1*^Cre^ (KP^fl^C) and *Kras*^G12D^; *Trp53*^R172H/+^; *Pdx1*^Cre^ (KPC) were used^19, 53^. Mice were monitored at least 3 times weekly. KP^fl^C and KPC mice were humanely killed upon detection of a palpable primary tumour, prior to the development of macroscopic metastasis, and age matched WT controls and KC mice were humanely killed at the same time.

### Tissue preparation for flow cytometry

Blood was collected in 2mM EDTA (ThermoFisher) coated syringes by cardiac puncture under terminal anaesthesia (sodium pentobarbital, i.p.) and red blood cells (RBC) were lysed. BM cells were harvested from the femur by flushing the bone through a 70µm filter with RPMI/5% (v/v) FBS/2mM EDTA. Lungs were excised and then minced with scissors before incubation in RPMI (ThermoFisher) with 10% foetal bovine serum (FBS, ThermoFisher) at 37°C for 20 minutes with agitation. Post-incubation, lungs were mashed through a 70µm filter. Spleens were excised and mashed through a 70µm filter. Livers and liver tumours were excised and then minced with scissors before incubation in RPMI/10% (v/v) FBS/2mM EDTA/1.2mg/mL Collagenase D (Roche)/1mg/mL DNase I (Roche) at 37°C for 20 minutes with agitation. Post-incubation livers were mashed through a 70µm followed by a 40µm filter. A section of small intestine was excised, flushed with RPMI/2% (v/v) FBS and minced with scissors before incubation in RPMI/75µg/mL DNase I/2mg/mL Collagenase D at 37°C for 30 minutes with agitation. Post-incubation, small intestines were mashed through a 70µm filter followed by a 40µm filter. Pancreatic tumours were excised and then minced with scissors before incubation in RPMI/75µg/mL DNase I/2mg/mL Collagenase D at 37°C for 30 minutes with agitation. Post-incubation, pancreatic tumours were mashed through a 70µm filter followed by a 40µm filter. Intestinal tumours were excised and then minced with scissors before incubation in RPMI/75µg/mL DNase I/2mg/mL Collagenase D at 37°C for 30 minutes with agitation. Post-incubation intestinal tumours were mashed through a 70µm filter followed by a 40µm filter.

### Extracellular protein staining for flow cytometry analysis

Single cell suspensions were stained with Live/Dead stain (LIVE/DEAD Fixable Near-IR or Yellow, Biolegend) and Fc-Receptor block was performed (using clone 93, Biolegend). Cell suspensions were incubated with directly conjugated fluorescent antibodies in PBS/1% (w/v) BSA (Merck)/0.05% (v/v) Sodium Azide (GBiosciences) (PBA) buffer at 4°C for 30 minutes. Biotinylated antibodies were labelled with fluorescently conjugated Streptavidin in PBA at 4°C for 30 minutes.

### Intracellular protein staining for flow cytometry analysis

Following extracellular protein staining, cells were fixed and permeabilized (Invitrogen) prior to staining with directly conjugated fluorescent antibodies at 4°C for 30-60 minutes. For Ki-67 staining, after extracellular staining cells were fixed with 2% PFA for 10 minutes at room temperature and washed with PBS. Cells were then re-suspended in fluorescently-conjugated Ki-67 antibody in PBS for 25 minutes at room temperature. Cells were washed with PBS prior to analysis.

### Flow cytometry acquisition and analysis

Count beads (CountBright, Life Technologies) were used to determine cell numbers. Compensation was performed using single stained beads (Ultracomp eBeads, ThermoFisher). Fluorescence minus one (FMO) controls were used to determine accurate gating. Acquisition was performed on a BDFortessa using FacsDiva software (BD Biosciences) and data analysed using FlowJo software (BD).

### Tissue preparation for Fluorescence Activated Cell Sorting (FACS)

For FACS of neutrophils for RNA-Seq analysis, PB, BM, SP and LV was collected as for flow cytometry. For *ex vivo* culture and adoptive transfer of neutrophils post-sorting, BM cells were harvested from the femur, tibia, radius and ulna dependent on the cell numbers required and collected into PBS/5% (v/v) FBS/1% (v/v) Penicillin/Streptomycin (ThermoFisher). All subsequent processing was undertaken in a biological safety level 2 tissue culture hood with sterile reagents. Bones were washed in 70% EtOH and flushed as for flow cytometry with RPMI/5% (v/v) FBS/2mM EDTA/1% (v/v) Penicillin/Streptomycin. BM cells were enriched for neutrophils using a mouse neutrophil negative enrichment kit (Biolegend).

### Extracellular protein staining for FACS

For RNA-Seq analysis, single cell suspensions were stained as for flow cytometry, however Live/Dead stain was not performed and instead 4’-6’-diamidino-2-phenylindole (DAPI, ThermoFisher) was added immediately prior to sorting for live/dead detection. For *ex vivo* culture and adoptive transfer of neutrophils post-sorting, single cell suspensions were stained as for flow cytometry but in sterile PBS/1% (v/v) FBS/1mM EDTA. For sorting, cells were re-suspended in PBS/1% (v/v) FBS/1% (v/v) Pen/Strep/5mM EDTA/25mM HEPES (ThermoFisher).

### FACS acquisition

Acquisition was performed on a BD FACSAria sorter situated within a biological safety level 2 enclosure at 4°C using a 100µm nozzle at the slowest acquisition speed and at a flow rate of approximately 2,000 events/second. Sorting strategy was optimised and determined to be ≥97% by re-running a small proportion of the sorted fractions on the cytometer. Sample processing, gating strategy and parameters for all sorting were kept consistent after optimisation. For RNA-Seq analysis, populations were sorted into 15mL Falcon tubes coated with PBS/10% (v/v) FBS and containing PBA buffer. For *ex* vivo culture and adoptive transfer experiments, neutrophil populations were sorted into 15mL Falcon tubes coated with RPMI/10% (v/v FBS) and containing RPMI/20% (v/v) FBS/5mM EDTA/25mM HEPES/1% (v/v) Pen/Strep.

### Antibodies and reagents for flow cytometry/FACS

The following antibodies were used: CD45 (clone 30-F11), CD11b (clone M1/70), Ly6G (clone 1A8), CD3e (clone 145-2C11), CD19 (clone 6D5), Nkp46 (clone 29A1.4), Ter119 (clone TER119) ICAM-1 (clone YN1/1.7.4), CD115 (clone AFS98), CD117 (clone 2B8), Sca-1 (clone D7), CCR6 (clone 29-2L17), CD48 (clone HM48-1), Ki-67 (clone 16A8), CD16/32 (clone 93), CD18 (clone M18/2), CXCR4 (clone L276F12), CXCR2 (clone SA045E1), Streptavidin (all Biolegend), Ly6B.2 (clone 7/4, Bio-Rad), CD133 (clone 13A4, eBioscience), CD49d (clone R1-2, ThermoFisher), CD101 (clone Moushi101, ThermoFisher), Siglec-F (clone E50-2440, BD Biosciences), CCR1 (clone 643854, Bio-Techne).

### Cell trace staining

Sorted BM Ly6G^Int^ and Ly6G^Hi^ neutrophil populations, and CD3^+^ T cells enriched from the spleen of naïve mice, were incubated for 15 minutes at 37°C with CellTrace CFDA-SE (ThermoFisher), CellTrace Yellow (ThermoFisher) or CellTrace Red (ThermoFisher) diluted in sterile PBS. Cells were washed in

### CSF-3-induced acute haematopoietic stress

Female C57BL/6 mice received a single dose of mouse recombinant CSF-3 (StemCell Technologies) at 0.25mg/kg in sterile PBS, or sterile PBS alone for control, by i.p. injection for the specified time-period at which point the mice were sacrificed.

### LPS-induced acute systemic inflammation

Mice received a single dose of LPS 0111:B4 from *Escheria coli* (Merck) at 1mg/kg in sterile PBS, or sterile PBS alone for control, by i.p. injection for the specified time-period at which point the mice were sacrificed.

### EdU incorporation assay

Female C57BL/6 mice received a single dose of the thymidine analogue EdU (ThermoFisher) at 50mg/kg in sterile PBS by i.p. injection for the specified time-period at which point the mice were sacrificed. Cells were isolated and labelled with directly conjugated fluorescent antibodies as for flow cytometry. After labelling, cells were fixed and stained for EdU following the manufacturer’s instruction and analysed by flow cytometry.

### Neutrophil adoptive transfer and lung extraction

Ly6G^Int^ and Ly6G^Hi^ neutrophil populations were FACS sorted from the BM of CSF-3 challenged mice and cell trace stained, as previously described. Post-cell trace staining, 1.5x10^6^ cells from each neutrophil population was mixed into a final volume of 100µL and intravenously (i.v.) injected into the tail vein of PBS control or LPS challenged mice. Mice were sacrificed and blood was collected as for flow cytometry. The right lung lobe was tied off, a small incision made to the trachea and a customized blunted 22G needle inserted. The left lung was inflated with 0.6mL PBS/2% (w/v) low melting point agarose (Merck) in PBS. The right lung lobe was excised for processing by flow cytometry. The inflated left lobe was excised and fixed in PBS/4% (v/v) formaldehyde (VWR) for 2 hours at 4°C.

### Fixed precision cut lung slices (PCLS)

Fixed lungs were sliced into 300µm thick sections on a vibrating microtome (Campden Ltd). Slices were stained with Hamster anti-CD31 ab (2H8, Abcam) in PBS/1% (w/v) bovine serum albumin (BSA, Sigma)/10% (v/v) normal goat serum (NGS, Sigma)/0.001% (v/v) sodium-azide (VWR). Hamster-IgG (ThermoFisher) were used to control for unspecific staining. Anti-Hamster Cy3 (Jackson) was used as a secondary antibody and DAPI to stain nuclei.

### Confocal imaging

PCLS prepared and stained as described above were imaged on a Zeiss LSM880 confocal microscope, processed in Zeiss Software before quantification and analysis using Imaris software (Bitplane, Oxford Instruments).

### Bulk RNA isolation, sequencing and normalization

All mice used for RNA isolation were humanely killed within the same period in the morning (between 07.00- 08.30) to minimise circadian-induced gene expression changes between samples. RNA was isolated from Ly6G^Int^, Ly6G^Hi^ or Ly6G^+^ neutrophil populations that had been FACS sorted from the BM, spleen and peripheral blood. Biological replicates for PBS and LPS bulk Ly6G^+^ neutrophil RNA-Seq are as follows: PBS: peripheral blood Ly6G^Hi^ n=4 mice; BM Ly6G^Int^ n=4 mice; BM Ly6G^Hi^ n=4 mice; spleen Ly6G^Int^ n=6 mice; spleen Ly6G^Hi^ n=7 mice; liver Ly6G^+^ n=7 mice. LPS: peripheral blood Ly6G^Int^ n=4 mice; peripheral blood Ly6G^Hi^ n=4 mice; BM Ly6G^Int^ n=3 mice; BM Ly6G^Hi^ n=3 mice; spleen Ly6G^Int^ n=4 mice; spleen Ly6G^Hi^ n=4 mice; liver Ly6G^+^ n=4 mice. Biological replicates for PDAC model bulk Ly6G^+^ neutrophil RNA-Seq are as follows: BM: WT Ly6G^Int^ n=10 mice; KP^fl^C Ly6G^Int^ n=5 mice; KPC Ly6G^Int^ n=5 mice; KPC Ly6G^Hi^ n=5 mice. Spleen: WT Ly6G^Int^ n=8 mice; KP^fl^C Ly6G^Int^ n=4 mice; KPC Ly6G^Int^ n=5 mice.

RNA was extracted from sorted populations using an RNeasy Micro Kit (Qiagen). RNA quality was evaluated using an Agilent Bioanalyser using RNA Pico chips. Libraries for cluster generation and DNA sequencing were prepared using a TaKaRa SMARTer Stranded Total RNA-Seq – Pico Input Mammalian kit v2.0 as per the manufacturer’s instructions. Quality and quantity of the DNA libraries was assessed on an Agilent 2200 Tapestation (D1000 screentape) and Qubit (Thermo Fisher Scientific) respectively. The libraries were run on the Illumina Next Seq 500 using the High Output 75 cycles kit (2x36 cycles, paired end reads, dual index). Illumina data were demultiplexed using bcl2fastq version 2.19.0.216, then adaptor sequences were removed using Cutadapt version 0.6.4. Next, quality checks on fastq files were performed using fastqc version 0.11.8. Paired-end reads were then aligned to the mouse genome version GRCh38.95 using HISAT2 version 2.1.0 with the –known-splicesite-infile and –rna-strandness options. Expression levels were determined using HTseq version 0.11.2 with the -s=reverse, --mode=intersection-nonempty and –type=exon options, then statistically analysed using R version 3.5.1. Differential gene expression analysis based on the negative binomial distribution was performed using DESeq2 version 1.22.2^54^.

### Identification of reliably expressed genes and DEGs from RNA-Seq

From normalized data, genes with reliable expression were identified as those with ≥ 2 reads/million in at least half of the replicates per neutrophil population. Genes not matching these conditions were excluded. DEGs were identified as those with a fold change ≥ 1.5 and P-value ≤ 0.05 between compared populations.

### Bulk proteomic analysis

Ly6G^Int^ and Ly6G^Hi^ neutrophil populations were sorted from the peripheral blood and BM of CSF-3 (i.p. 0.25 mg/kg, 24 hours) administered mice. BM Ly6G^Int^ n=3, and peripheral blood and BM Ly6G^Hi^ n=3 replicates. Each replicate was composed of FACS isolated Ly6G^Int^ and Ly6G^Hi^ neutrophils pooled from n=4 mice. Sorted neutrophil populations were pelleted, snap frozen in liquid nitrogen and stored at -70°C. Sorted neutrophil populations from 4 mice per replica were thawed and pooled. Samples were lysed by boiling with sodium dodecyl sulfate (SDS), sonicated and centrifuged for debris removal. Proteins obtained in the supernatants were precipitated in two steps using 24% and 10% solution of trichloroacetic acid. In both steps, pellets were incubated at 4°C for 10 min and centrifuged at 18,000 g for 5 min. Supernatants were carefully aspirated and pellets were washed with water until the supernatant reached neutral pH. Pellets were reconstituted in… and digested, first using Endoproteinase Lys-C (ratio 1:33 enzyme:lysate, Alpha Laboratories) for 1 h at room temperature, then with trypsin (ratio 1:33 enzyme:lysate, Promega) overnight at 35°C. *Off-line HPLC fractionation:* TMT-labelled peptides were fractionated using high pH reverse phase chromatography on a C18 column (150 × 2.1 mm i.d. - Kinetex EVO (5 μm, 100 Å)) on a HPLC system (Agilent, LC 1260 Infinity II, Agilent). A two-step gradient was applied, from 1–28% B in 42 min, then from 28–46% B in 13 min. The column eluate was collected into 9 fractions subsequently submitted to MS analysis. *UHPLC-MS/MS analysis:* Fractionated tryptic digests were separated by nanoscale C18 reverse-phase liquid chromatography using an EASY-nLC II 1200 (Thermo Fisher Scientific) coupled to an Orbitrap Fusion Lumos mass spectrometer (Thermo Fisher Scientific). Samples were loaded into a 50 cm fused silica emitter (New Objective) packed in-house with ReproSil-Pur C18-AQ, 1.9 μm resin (Dr Maisch GmbH). The emitter was kept at 50 °C by means of a column oven (Sonation) integrated into the nanoelectrospray ion source (Thermo Fisher Scientific). Elution was carried out at a flow rate of 300 nl/min using a binary gradient with buffer A (2% acetonitrile) and B (80% acetonitrile), both containing 0.1% formic acid. Peptides from each fraction were eluted at using three different binary gradients optimised for different set of fractions as described previously^55^. An Active Background Ion Reduction Device (ESI Source Solutions) was used to decrease air contaminants signal level. Data were acquired using Xcalibur software (Thermo Fisher Scientific) on an Orbitrap Fusion Lumos mass spectrometer (Thermo Fisher Scientific). The mass spectrometer was operated in positive ion mode and used in data-dependent acquisition (DDA). A full scan was acquired over mass range of 350-1400 m/z, with 60,000 resolution at 200 m/z, for a maximum injection time of 20 ms and with a target value of 9e5 ions with “monoisotopic precursor selection” set to “Peptide” mode. Higher energy collisional dissociation fragmentation was performed on the 15 most intense ions, for a maximum injection time of 150 ms, or a target value of 500,000 ions. Peptide fragments were analysed in the Orbitrap at 50,000 resolution. *Proteomic Data analysis:* The MS Raw data were processed with MaxQuant software^56^ version 1.6.3.3 and searched with Andromeda search engine^57^, querying UniProt^58^ *Mus musculus* (12/08/2018; 25,198 entries). First and main searches were carried out with precursor mass tolerances of 20 ppm and 4.5 ppm, respectively, and MS/MS tolerance of 20 ppm. The minimum peptide length was set to six amino acids and specificity for trypsin cleavage was required, allowing up to two missed cleavage sites. MaxQuant was set to quantify on “Reporter ion MS2”, and TMT10plex was chose as Isobaric label. Interference between TMT channels were corrected by MaxQuant using the correction factors provided by the manufacturer. The “Filter by PIF” option was activated and a “Reporter ion tolerance” of 0.003 Da was used. Methionine oxidation and N-terminal acetylation were specified as variable modifications, and Cysteine carbamidomethylation was set as fixed modification. The peptide and protein false discovery rate (FDR) was set to 1 %. MaxQuant output was further processed and analysed using Perseus software version 1.6.2.3. Protein quantification was done using the ProteinGroups.txt file, Reverse and Potential Contaminant proteins were removed, and at least one uniquely assigned peptide and a minimum ratio count of 2 were required for a protein to be quantified. To identify regulated proteins, ANOVA “Multiple sample test” in Perseus was used with a truncation threshold value of 0.001 (Permutation based FDR) and S0=0. The test was applied to the 3 groups considered. For data visualisation, proteins that passed the ANOVA test were z-score normalised. Hierarchical clustering analysis was performed using “Euclidian distance” for both rows and columns.

### Splenic T-cell enrichment and *ex vivo* suppression assays

T cells were enriched from single cell suspensions of splenocytes isolated from naïve C57BL/6 mice as for flow cytometry using a mouse CD3 T cell negative selection kit (Biolegend). Enriched CD3^+^ T cells were cell trace stained and cultured at 1x10^6^ cells/mL and stimulated with mouse T-activator anti-CD3/CD28 Dynabeads (ThermoFisher) at a 1:1 ratio. T cells were incubated with Ly6G^Int^ or Ly6G^Hi^ neutrophil populations sorted from the BM of naïve or CSF-3 challenged mice at varying ratios for 48 hours at 37°C, 5% CO2. Post-incubation cells were stained with a live/dead stain, labelled with anti-CD4 and anti-CD8 fluorescently conjugated antibodies and analysed by flow cytometry.

### Neutrophil *ex vivo* culture

Ly6G^Int^ and Ly6G^Hi^ neutrophil populations were sorted from the BM of naïve C57BL/6 mice. 1.75x10^3^ cells were loaded into individual well of a 96 well v-bottom plate in RPMI/5% (v/v) FBS/2mM EDTA/1% (v/v) Penicillin/Streptomycin/1% (v/v) L-Glutamine (ThermoFisher)/0.1% (v/v) β-Mercaptoethanol (ThermoFisher). Cells were incubated for the required time at 37°C, 5% CO2. Post-incubation total cell numbers were manually counted using a haemocytometer (Hausser Scientific) and stained for flow cytometry analysis.

### Data availability

The GEO accession code for RNA-Seq data is GSE180824. The mass spectrometry proteomics data have been deposited to the ProteomeXchange Consortium via the PRIDE partner repository^59^ with the dataset identifier PXD026708. All other data are available at reasonable request by contacting the corresponding author.

### Quantification and statistical analysis

Statistical analyses were performed using GraphPad Prism software (v8.0 GraphPad software, USA). Statistical tests used are as specified in figure legends.

### Author Contributions

JBGM and LMC conceived the study. JBGM, AJM, TJ, RJ, XLR, JS, WC, AH, KG, XC, and SL performed experiments. JBGM, AJM, XLR, WC, AH, KG, FF, SL, JV, SZ and LMC analysed and interpreted data. AJM, TJ, RJ, XLR, GJG, KDF, DJM, CWS, JCN, TGB, SZ, JM, OJS advised on and provided mouse models. JBGM and LMC wrote the manuscript. All authors reviewed and edited the manuscript.

## Acknowledgements

**General:** We would like to thank the core services and advanced technologies at the National Heart and Lung Institute and the CRUK Beatson Institute with particular thanks to the Biological Services Unit, Molecular Services, Bioinformatics and the Beatson Advanced Imaging Resource with particular thanks to Tom Gilbey for cell sorting. We would like to thank the members of the Leukocyte Dynamics Group, Seth Coffelt and Catherine Winchester for critical reading of the manuscript. We would also like to thank Sara Rankin and Clare Lloyd for helpful discussions during the project. Figure artwork made use of images from https://smart.servier.com/ Servier Medical Art, licensed under a Creative Commons Attribution 3.0 Unported Licence.

## Funding

JBGM was supported by an Imperial College London NHLI Foundation studentship, MRC grant (MR/M01245X/1) and CRUK core funding (A17196 and A31287). LMC (CRUK A23983), OJS (CRUK A21339), JCN (CRUK A18277), JM (CRUK A29996) and SZ (CRUK A29800) were supported by CRUK core funding at Cancer Research UK Beatson Institute. TGB was funded by Wellcome (WT107492Z) and the CRUK HUNTER Accelerator Award (CRUK A26813). D.A.M., O.J.S., T.G.B and J.B.G.M are supported by a CRUK program grant (C18342/A23390).

## Competing interests

DAM is a paid director and shareholder of Fibrofind limited. DJM receives research funding from Puma Bio-technology and Merck. OJS and TGB receive research funding from AstraZeneca.

## Notes

### Summary of Updates

Accession code and funding information updated

