## Extended Data Supplement for "Maturation, developmental site, and pathology dictate murine neutrophil function"

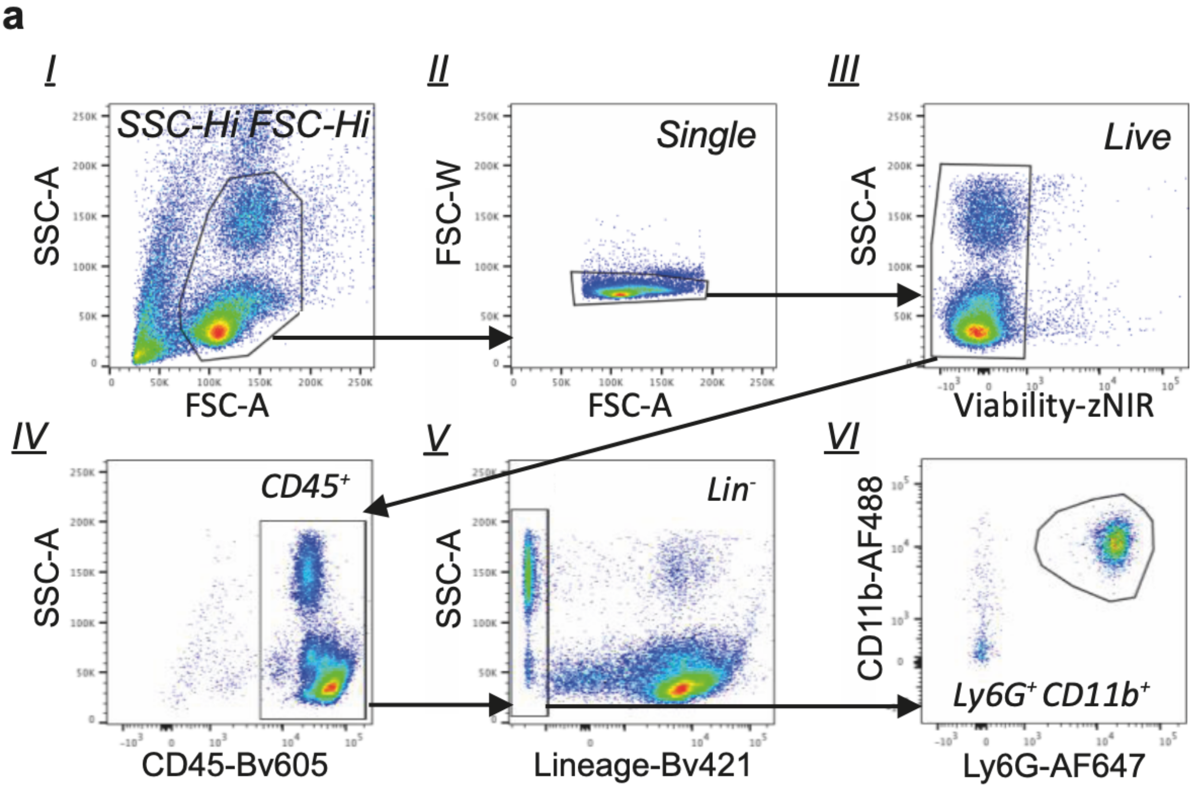


**Extended Data Fig. 1. Gating strategy for identification of Ly6G⁺CD11b⁺ neutrophils. a**, $SSC^{Hi}FSC^{Hi}$ (I), single cells (II), live cells (III), $CD45^{+}$ (IV), $Lineage^{-}$ (CD3, CD19, Ter119, CD115, Nkp46, Siglec-F and F4/80; (V)) and $Ly6G^{+}CD11b^{+}$ (VI).

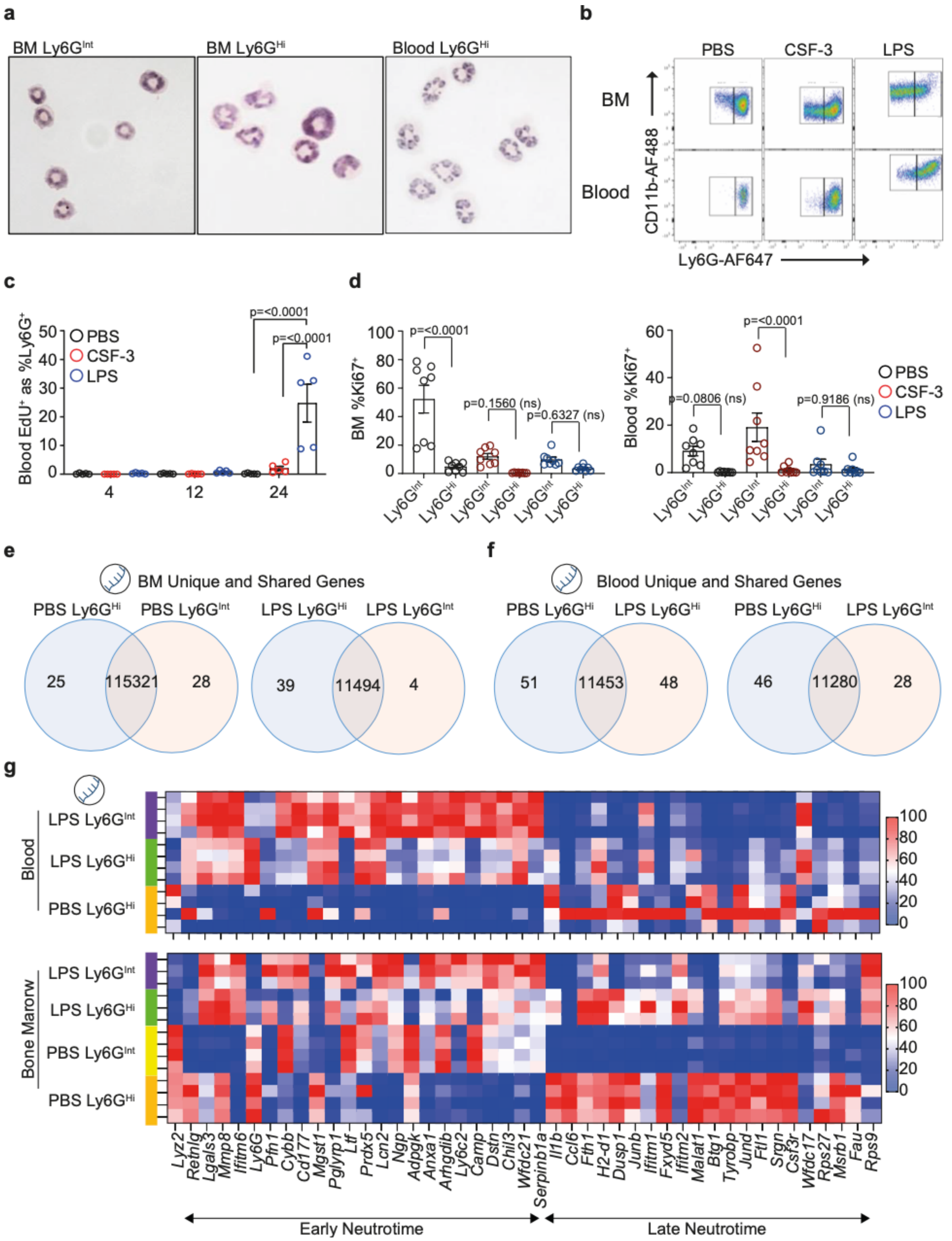


**Extended Data Fig. 2. Intermediate Ly6G expressing (Ly6G$^{Int}$) neutrophils are immature. a**, H&E stained cytospins of FACS isolated BM Ly6G$^{Int}$, BM Ly6G$^{Hi}$ and peripheral blood Ly6G$^+$ neutrophils from naïve mice. Data are representative of n=5 mice. **b**, Flow cytometry plots for Ly6G$^+$CD11b$^+$ neutrophils in PBS-control, CSF-3- and LPS-challenged mice. Data representative of: PBS n=11 mice; CSF-3 n=9 mice; LPS n=5 mice. **c**, Quantification of EdU$^+$ as %Ly6G$^+$ neutrophils in the peripheral blood. PBS n=5 mice; CSF-3 n=5 mice, LPS n=5 mice. Data were analysed by 2way ANOVA with Tukey's multiple comparisons test. **d**, Quantification of Ki-67$^+$ as %Ly6G$^{Int}$ and Ly6G$^{Hi}$ neutrophils in the BM (left) and peripheral blood (right). Ly6G$^{Int}$ and Ly6G$^{Hi}$ neutrophils: PBS n=8 mice;

CSF-3 n=8 mice; LPS n=8 mice. Data were analysed by 2way ANOVA with Sidak's multiple comparisons test. **e,** Venn diagrams for genes with unique and shared expression between $Ly6G^{Int}$ and $Ly6G^{Hi}$ neutrophils isolated from the BM of PBS-control (left) and LPS-challenged (right) mice. Data from transcriptomic analysis. **f**, Venn diagrams for genes with unique and shared expression between; PBS-control $Ly6G^{Hi}$ and LPS-challenged $Ly6G^{Hi}$ (left) and LPS-challenged $Ly6G^{Hi}$ and $Ly6G^{Int}$ neutrophils (right). Data from transcriptomic analysis. **g**, Heatmap showing row scaled expression for genes associated with early and late stages of neutrophil development (neutrotime, identified by Grieshaber-Bouyer *et al.,* 2020) for $Ly6G^{Int}$ and $Ly6G^{Hi}$ neutrophils in the peripheral blood and BM of PBS-control and LPS-challenged mice. Data from transcriptomic analysis.

Bulk $Ly6G^{Int}$ and $Ly6G^{Hi}$ neutrophil RNA-Seq data in Extended Data Fig. 2e-g. PBS BM: $Ly6G^{Int}$ n=4 mice; $Ly6G^{Hi}$ n=4 mice. LPS-BM: $Ly6G^{Int}$ n=3 mice; $Ly6G^{Hi}$ n=3 mice. PBS peripheral blood: $Ly6G^{Hi}$ n=4 mice. LPS peripheral blood: $Ly6G^{Int}$ n=4 mice; $Ly6G^{Hi}$ n=4 mice. Dots in Extended Data Fig. 2c, d represent individual mice. Error bars in Extended Data Fig. 2c, d represent mean±SEM.

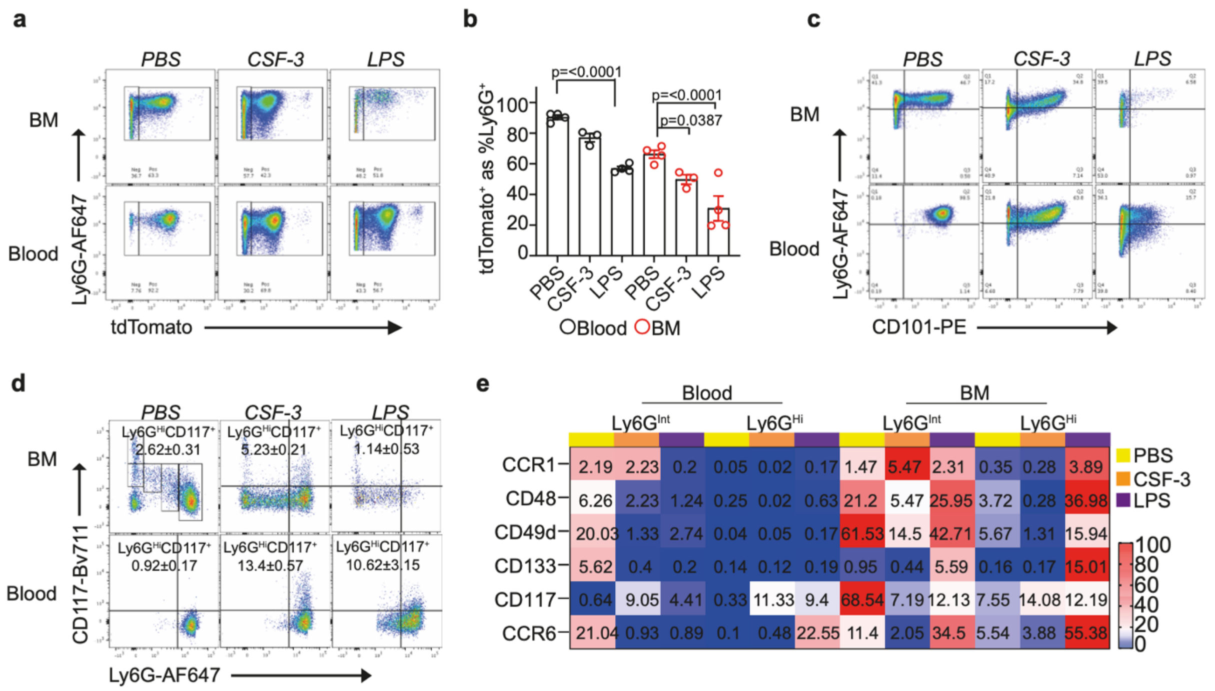


**Extended Data Fig. 3. Haematopoietic stress reduces the accuracy of alternative methods for identification of neutrophil maturity. a**, Flow cytometry plots for TdTomato expression by Ly6G⁺ neutrophils in PBS-control, CSF-3- and LPS-challenged mice. BM representative of: PBS n=4 mice; CSF-3 n=3 mice; LPS n=4 mice. Peripheral blood representative of: PBS n=4 mice; CSF-3 n=3 mice; LPS n=4 mice. **b**, Quantification of TdTomato⁺ as %Ly6G⁺ neutrophils. Peripheral blood Ly6G⁺ neutrophils: PBS n=4 mice; CSF-3 n=3 mice; LPS n=4 mice. BM Ly6G⁺ neutrophils: PBS n=4 mice; CSF-3 n=3 mice; LPS n=4 mice. Data were analysed by 2way ANOVA with Tukey's multiple comparisons test. **c**, Flow cytometry plots for CD101 expression by Ly6G⁺ neutrophils in PBS-control, CSF-3- and LPS-challenged mice. BM representative of: PBS n=5 mice; CSF-3 n=5 mice; LPS n=5 mice. Peripheral blood representative of: PBS n=5 mice; CSF-3 n=5 mice; LPS n=5 mice. **d**, Flow cytometry plots for CD117 expression by Ly6G⁺ neutrophils in PBS-control, CSF-3- and LPS-challenged mice. Data are presented as mean±standard deviation. BM representative of: PBS n=3 mice; CSF-3 n=3 mice; LPS n=3 mice. Peripheral blood representative of: PBS n=3 mice; CSF-3 n=3 mice; LPS n=3 mice. **e**, Heatmap showing row scaled CCR1⁺, CD48⁺, CD49d⁺, CD133⁺, CD117⁺ and CCR6⁺ as %Ly6G$^{Int}$ and Ly6G$^{Hi}$ neutrophils isolated from the peripheral blood and BM of PBS-control, CSF-3- and LPS-challenged mice. Data from flow cytometry analysis. Peripheral blood Ly6G$^{Int}$ and Ly6G$^{Hi}$ neutrophils: CCR1 PBS n=7 mice, CSF-3 n=4 mice, LPS n=4 mice; CD48 PBS n=7 mice, CSF-3 n=4 mice; LPS n=4 mice; CD49d PBS n=5 mice, CSF-3 n=7 mice, LPS n=7 mice; CD133 PBS n=4 mice, CSF-3 n=4 mice, LPS n=4 mice; CD117 PBS n=5 mice, CSF-3 n= 7 mice, LPS n=7 mice; CCR6 PBS n=4 mice, CSF-3 n=4 mice, LPS n=4 mice. BM Ly6G$^{Int}$ and Ly6G$^{Hi}$ neutrophils: CCR1 PBS n=7 mice, CSF-3 n=4 mice, LPS n=4 mice; CD48 PBS n=7 mice, CSF-3 n=4 mice; LPS n=4 mice; CD49d PBS n=4 mice, CSF-3 n=7 mice, LPS n=7 mice; CD133 PBS n=4 mice, CSF-3 n=4 mice, LPS n=4 mice; CD117 PBS n=5 mice, CSF-3 n= 7 mice, LPS n=7 mice; CCR6 PBS n=4 mice, CSF-3 n=4 mice, LPS n=4 mice.

Dots in Dots in Extended Data Fig. 3b represent individual mice. Error bars in Extended Data Fig. 3b represent mean±SEM.

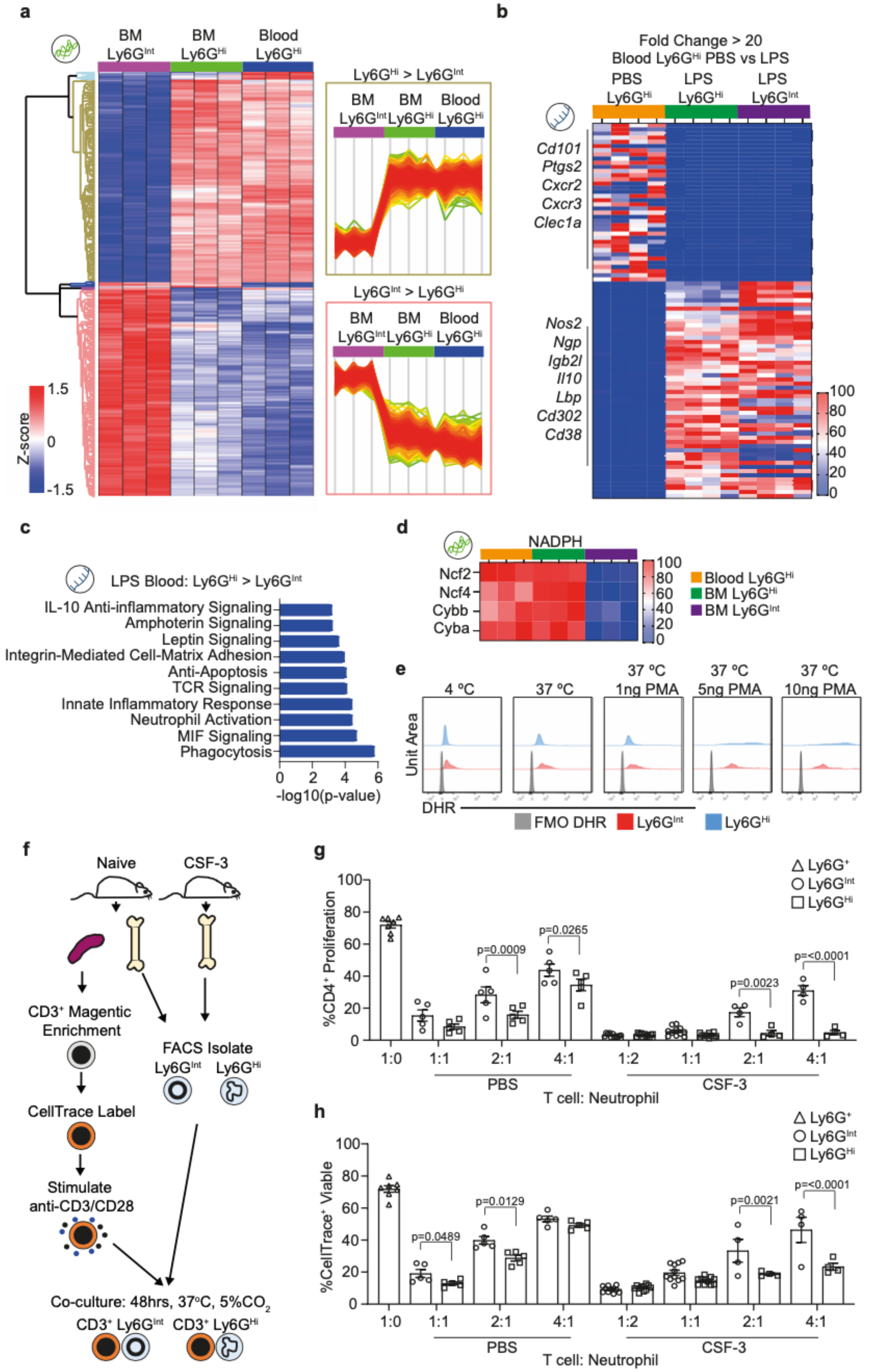
a
BM Ly6GInt
BM Ly6GHi
Blood Ly6GHi
Ly6GHi > Ly6GInt
Ly6GInt > Ly6GHi
Z-score
1.5
0
-1.5
b
Fold Change > 20
Blood Ly6GHi PBS vs LPS
PBS Ly6GHi
LPS Ly6GHi
LPS Ly6GInt
Cd101
Ptgs2
Cxcr2
Cxcr3
Clec1a
Nos2
Ngp
Igb2l
Il10
Lbp
Cd302
Cd38
c
LPS Blood: Ly6GHi > Ly6GInt
IL-10 Anti-inflammatory Signaling
Amphoterin Signaling
Leptin Signaling
Integrin-Mediated Cell-Matrix Adhesion
Anti-Apoptosis
TCR Signaling
Innate Inflammatory Response
Neutrophil Activation
MIF Signaling
Phagocytosis
-log10(p-value)
d
NADPH
Ncf2
Ncf4
Cybb
Cyba
Blood Ly6GHi
BM Ly6GHi
BM Ly6GInt
e
4 °C
37 °C
37 °C 1ng PMA
37 °C 5ng PMA
37 °C 10ng PMA
Unit Area
DHR
FMO DHR
Ly6GInt
Ly6GHi
f
Naive
CSF-3
CD3+ Magentic Enrichment
CellTrace Label
Stimulate anti-CD3/CD28
FACS Isolate
Ly6GInt
Ly6GHi
Co-culture: 48hrs, 37°C, 5%CO2
CD3+ Ly6GInt
CD3+ Ly6GHi
g
%CD4+ Proliferation
Ly6G+
Ly6GInt
Ly6GHi
p=0.0009
p=0.0265
p=0.0023
p=<0.0001
1:0
1:1
2:1
4:1
1:2
PBS
CSF-3
T cell: Neutrophil
h
%CellTrace+ Viable
p=0.0489
p=0.0129
p=0.0021
p=<0.0001

**Extended Data Fig. 4. Ly6G$^{Int}$ display transcriptomic, proteomic and functional differences to Ly6G$^{Hi}$ neutrophils. a,** Heatmap showing row scaled protein abundance between Ly6G$^{Int}$ and Ly6G$^{Hi}$ neutrophils isolated from the peripheral blood and BM of CSF-3-challenged mice. Data from proteomic analysis. **b**, Heatmap showing row scaled expression for genes with >20 fold change between peripheral blood Ly6G$^{Hi}$ neutrophils isolated from PBS-control and LPS-challenged mice. Data from transcriptomic analysis. **c**, Top 10 most enriched Process Networks from DEGs with increased expression in peripheral blood Ly6G$^{Hi}$ compared to Ly6G$^{Int}$ neutrophils from LPS-challenged mice. Data from transcriptomic analysis. **d**, Heatmap showing row scaled protein abundance for NADPH components between Ly6G$^{Int}$ and Ly6G$^{Hi}$ neutrophils isolated from the peripheral blood and BM of CSF-3 challenged mice. Data from proteomic analysis. **e**, Histograms showing intracellular-DHR-123 fluorescence for *ex vivo* PMA stimulated Ly6G$^{Int}$ and Ly6G$^{Hi}$ neutrophils isolated from the BM of naïve mice. Data representative of Ly6G$^{Int}$ and Ly6G$^{Hi}$ neutrophils n=6 mice for each condition. **f**, Schematic for *ex vivo* neutrophil suppression assays. **g**, Quantification of %CD4$^+$ T cell proliferation when cultured *ex vivo* with Ly6G$^{Int}$ or Ly6G$^{Hi}$ neutrophils isolated from the BM of PBS-control or CSF-3-challenged mice, at different T cell: neutrophil ratios. Data from flow cytometry analysis. Ly6G$^+$ neutrophils 1:0 ratio n=7 mice. Ly6G$^{Int}$ and Ly6G$^{Hi}$ neutrophils: PBS 1:1, 2:1 and 4:1 ratios n=5 mice; CSF-3 1:2 and 1:1 ratios n=11 mice; CSF-3 2:1 and 4:1 ratios n=4 mice. Data were analysed by 2way ANOVA with Sidak's multiple comparisons test. **h**, Quantification of %Viable CellTrace$^+$ CD3$^+$ T cells when cultured *ex vivo* with Ly6G$^{nt}$ or Ly6G$^{Hi}$ neutrophils isolated from the BM of PBS-control or CSF-3-challenged mice, at different T cell: neutrophil ratios. Data from flow cytometry analysis. Ly6G$^+$ neutrophils 1:0 ratio n=7 mice. Ly6G$^{Int}$ and Ly6G$^{Hi}$ neutrophils: PBS 1:1, 2:1 and 4:1 ratios n=5 mice; CSF-3 1:2 and 1:1 ratios n=11 mice; CSF-3 2:1 and 4:1 ratios n=4 mice. Data were analysed by 2way ANOVA with Sidak's multiple comparisons test.

Bulk Ly6G$^{Int}$ and Ly6G$^{Hi}$ neutrophil RNA-Seq data in Extended Data Fig. 4b, c. PBS BM: Ly6G$^{Int}$ n=4 mice; Ly6G$^{Hi}$ n=4 mice. LPS-BM: Ly6G$^{Int}$ n=3 mice; Ly6G$^{Hi}$ n=3 mice. PBS peripheral blood: Ly6G$^{Hi}$ n=4 mice. LPS peripheral blood: Ly6G$^{Int}$ n=4 mice; Ly6G$^{Hi}$ n=4 mice. Bulk Ly6G$^{Int}$ and Ly6G$^{Hi}$ neutrophil proteomic data in Fig. 2l. CSF-3 BM Ly6G$^{Int}$ n=3, and peripheral blood and BM Ly6G$^{Hi}$ n=3 replicates. Each replicate composed of FACS isolated Ly6G$^{Int}$ and Ly6G$^{Hi}$ neutrophils pooled from n=4 mice. Dots in Dots in Extended Data Fig. 4g, h represent individual mice. Error bars in Extended Data Fig. 4g, h represent mean±SEM.

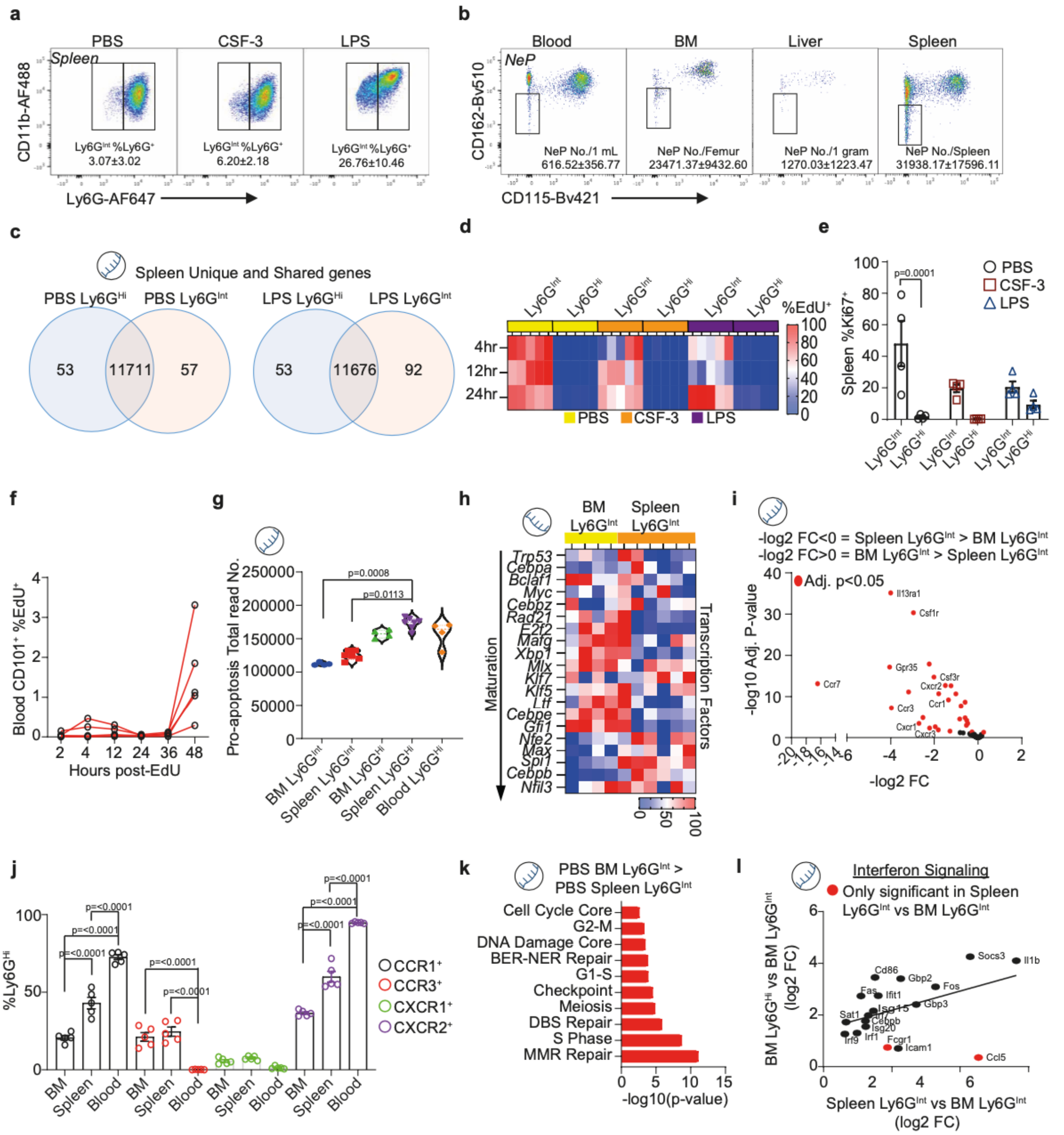


**Extended Data Fig. 5. Splenic Ly6G$^{Int}$ are immature and distinct from their BM counterparts. a,** Flow cytometry plots for Ly6G$^{Int}$CD11b$^{+}$ neutrophils in the spleens of PBS-control, CSF-3- and LPS-challenged mice. Data are presented as mean±standard deviation. Data representative of: PBS n=7; CSF-3 n=7; LPS n=7. **b**, Flow cytometry plots for the unipotent neutrophil progenitor, termed NeP (characterised by Zhu *et al.,* 2018), in the peripheral blood, BM, liver and spleen of naïve mice. Data are presented as mean±standard deviation. Data are representative of: peripheral blood n=5 mice; BM n=5 mice; liver n=5 mice; spleen n=5 mice. **c**, Venn diagrams for genes with unique and shared expression between Ly6G$^{Hi}$ and Ly6G$^{Int}$ neutrophils isolated from the spleen of PBS-control (left) and LPS-challenged (right) mice. Data from transcriptomic analysis. **d**, Heatmap showing row-scaled EdU$^{+}$ as %Ly6G$^{Int}$ and Ly6G$^{Hi}$ neutrophils isolated from the spleen at 4, 12 and 24 hours post-EdU injection. PBS 4hr, 12hr and 24hr Ly6G$^{Int}$ and Ly6G$^{Hi}$ n=5 mice; CSF-3 4hr, 12hr and 24hr Ly6G$^{Int}$ and Ly6G$^{Hi}$ n=5 mice; LPS 4hr, 12hr and 24hr Ly6G$^{Int}$ and Ly6G$^{Hi}$ n=5 mice. **e**, Quantification of Ki-67$^{+}$ as %Ly6G$^{Int}$ and Ly6G$^{Hi}$ neutrophils in the spleen. PBS Ly6G$^{Int}$ and Ly6G$^{Hi}$ n=4 mice; CSF-3 Ly6G$^{Int}$ and Ly6G$^{Hi}$ n=4 mice; LPS Ly6G$^{Int}$ and Ly6G$^{Hi}$ n=4 mice. Data were analysed by 2way ANOVA with Sidak's multiple comparisons test. **f**, Quantification of CD101$^{+}$ as %EdU$^{+}$ neutrophils in the peripheral blood of naïve mice. At all time-points n=5 mice. **g**, Quantification of total combined read no. for genes associated with pro-apoptosis (GO:0043065). Data from transcriptomic analysis. **h**, Heatmap showing column scaled expression for transcription factors associated with neutrophil maturation. Data from transcriptomic analysis. **i**, Volcano plot for genes associated with leukocyte chemotaxis (GO:0030595) comparing BM Ly6G$^{Int}$ and spleen Ly6G$^{Int}$ neutrophils isolated from naïve mice. Data from transcriptomic analysis. Fold change = FC. **j**, Quantification of CCR1$^{+}$, CCR3$^{+}$, CXCR1$^{+}$ and CXCR2$^{+}$ as %Ly6G$^{Hi}$ neutrophils in the BM, spleen and peripheral blood of naïve mice. CCR1$^{+}$ BM, spleen, peripheral blood n=5 mice; CCR3$^{+}$ BM, spleen, peripheral blood n=5 mice; CXCR1$^{+}$ BM, spleen, peripheral blood n=5 mice; CXCR2$^{+}$ BM, spleen, peripheral blood n=5 mice. Data were analysed by 2way ANOVA with Tukey's multiple comparisons test. **k**, Top 10 most enriched Process Networks from DEGs with increased expression in BM Ly6G$^{Int}$ compared to spleen Ly6G$^{Int}$ neutrophils from PBS-control mice. Data from transcriptomic analysis. **l**, Correlation for DEGs increased in PBS-control spleen Ly6G$^{Int}$ compared to BM Ly6G$^{Int}$ with DEGs increased in PBS-control BM Ly6G$^{Hi}$ compared to BM Ly6G$^{Int}$, associated with the process network Inflammation Interferon Signaling. Data from transcriptomic analysis. Fold change = FC.

Bulk Ly6G$^{Int}$ and Ly6G$^{Hi}$ neutrophil RNA-Seq data in Extended Data Fig. 5c, g-i, k, l. PBS BM: Ly6G$^{Int}$ n=4 mice; Ly6G$^{Hi}$ n=4 mice. LPS-BM: Ly6G$^{Int}$ n=3 mice; Ly6G$^{Hi}$ n=3 mice. PBS peripheral blood: Ly6G$^{Hi}$ n=4 mice. LPS peripheral blood: Ly6G$^{Int}$ n=4 mice; Ly6G$^{Hi}$ n=4 mice. Dots in Extended Data Fig. 5e-g, j represent individual mice. Error bars in Extended Data Fig. 5e, j represent mean±SEM.

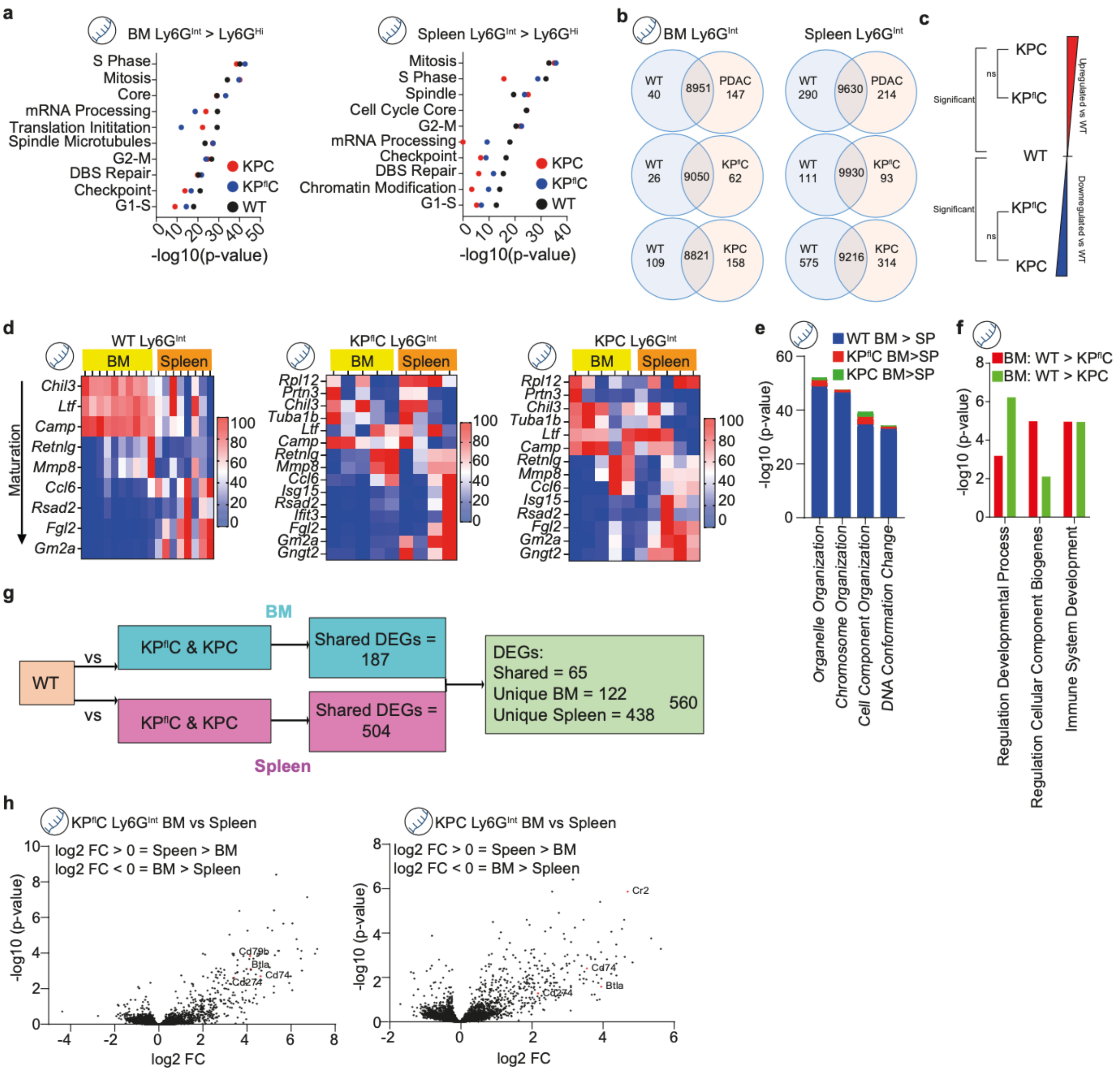


**Extended Data Fig. 6. Ly6G$^{Int}$ neutrophils maintain their immature status and tissue specific phenotypes in tumour bearing mice. a**, DEGs associated with cell cycle regulation with increased expression in BM Ly6G$^{Int}$ compared to Ly6G$^{Hi}$ (left) and spleen Ly6G$^{Int}$ compared to Ly6G$^{Hi}$ (right) in WT, KP$^{fl}$C and KPC mice, shown as -Log10(p-value). Data from transcriptomic analysis. **b**, Venn diagrams for DEGs shared by KP$^{fl}$C and KPC (PDAC), unique to KP$^{fl}$C and unique to KPC compared to WT for BM Ly6G$^{Int}$ (left) and spleen Ly6G$^{Int}$ neutrophils (right). Data from transcriptomic analysis. **c**, Schematic showing proposed gene expression for Ly6G$^{Int}$ neutrophils from KP$^{fl}$C and KPC mice compared to WT. **d**, Heatmap showing row scaled expression for genes associated with neutrophil maturation (identified by Xie *et al.,* 2020) for BM Ly6G$^{Int}$ and spleen Ly6G$^{Int}$ neutrophils from WT (left), KP$^{fl}$C (middle) and KPC mice (right). Data from transcriptomic analysis. **e**, Enrichment analysis for GO Processes associated with neutrophil development increased in BM Ly6G$^{Int}$ compared to spleen Ly6G$^{Int}$ neutrophils from WT, KP$^{fl}$C and KPC mice. Data from transcriptomic analysis. **f**, Enrichment analysis for GO Processes associated with neutrophil development increased in WT BM Ly6G$^{Int}$ compared to KP$^{fl}$C and KPC BM Ly6G$^{Int}$ neutrophils. Data from transcriptomic analysis. **g**, Schematic for identification and comparison of transcriptome changes identified in PDAC bearing compared to WT mice in BM and spleen Ly6G$^{Int}$ neutrophils. **h**, Volcano plot for 560 DEGs identified between BM and spleen Ly6G$^{Int}$ shared by KP$^{fl}$C (left) and KPC (right) mice compared to WT. Here, 122 DEGS are increased in BM > spleen Ly6G$^{Int}$ and 438 DEGs are increased in spleen > BM Ly6G$^{Int}$. Data from transcriptomic analysis. Fold change = FC.

Bulk Ly6G$^{Int}$ and Ly6G$^{Hi}$ neutrophil RNA-Seq data in Fig. 5b-f. BM: WT Ly6G$^{Int}$ n=10 mice; WT Ly6G$^{Hi}$ n=10 mice; KP$^{fl}$C Ly6G$^{Int}$ n=5 mice; KP$^{fl}$C Ly6G$^{Hi}$ n=5 mice; KPC Ly6G$^{Int}$ n=5 mice; KPC Ly6G$^{Hi}$ n=5 mice. Spleen: WT Ly6G$^{Int}$ n=8 mice; WT Ly6G$^{Hi}$ n=10 mice; KP$^{fl}$C Ly6G$^{Int}$ n=4 mice; KP$^{fl}$C Ly6G$^{Hi}$ n=5 mice; KPC Ly6G$^{Int}$ n=5 mice; KPC Ly6G$^{Hi}$ n=5 mice.

**Tables:**

| Peripheral blood vs peripheral blood | | | DEGs | Total DEGs |
|---|---|---|---|---|
| PBS BL Ly6GHi | > | LPS BL Ly6GHi | 2634 | 4374 |
| LPS BL Ly6GHi | > | PBS BL Ly6GHi | 1740 | |
| LPS BL Ly6GHi | > | LPS BL Ly6GInt | 567 | 1147 |
| LPS BL Ly6GInt | > | LPS BL Ly6GHi | 580 | |
| BM vs BM | | | DEGs | Total DEGs |
| PBS BM Ly6GHi | > | PBS BM Ly6GInt | 2004 | 5915 |
| PBS BM Ly6GInt | > | PBS BM Ly6GHi | 3911 | |
| LPS BM Ly6GHi | > | LPS BM Ly6GInt | 353 | 533 |
| LPS BM Ly6GInt | > | LPS BM Ly6GHi | 180 | |
| PBS BM Ly6GHi | > | LPS BM Ly6GHi | 1421 | 3672 |
| LPS BM Ly6GHi | > | PBS BM Ly6GHi | 2251 | |
| PBS BM Ly6GInt | > | LPS BM Ly6GInt | 1699 | 2786 |
| LPS BM Ly6GInt | > | PBS BM Ly6GInt | 1087 | |
| Spleen vs spleen | | | DEGs | Total DEGs |
| PBS SP Ly6GHi | > | PBS SP Ly6GInt | 2654 | 6896 |
| PBS SP Ly6GInt | > | PBS SP Ly6GHi | 4242 | |
| LPS SP Ly6GHi | > | LPS SP Ly6GInt | 1509 | 2717 |
| LPS SP Ly6GInt | > | LPS SP Ly6GHi | 1208 | |
| PBS SP Ly6GHi | > | LPS SP Ly6GHi | 2069 | 5448 |
| LPS SP Ly6GHi | > | PBS SP Ly6GHi | 3379 | |
| PBS SP Ly6GInt | > | LPS SP Ly6GInt | 2673 | 4000 |
| LPS SP Ly6GInt | > | PBS SP Ly6GInt | 1327 | |

**Table 1.** DEGs identified by p-value≤0.05 and fold change≥1.5 from comparisons of Ly6G$^{Hi}$ and Ly6G$^{Int}$ neutrophil populations isolated from the peripheral blood (BL), bone marrow (BM) and spleen (SP) of PBS-control and LPS-challenged mice. Data from transcriptomic analysis.

| Ly6G$^{Int}$ Comparisons: Blood, BM, Spleen | | | DEGs | Total DEGs |
|---|---|---|---|---|
| PBS BM Ly6GInt | > | PBS SP Ly6GInt | 93 | 968 |
| PBS SP Ly6GInt | > | PBS BM Ly6GInt | 875 | |
| LPS BM Ly6GInt | > | LPS SP Ly6GInt | 112 | 684 |
| LPS SP Ly6GInt | > | LPS BM Ly6GInt | 572 | |
| LPS SP Ly6GInt | > | LPS PB Ly6GInt | 18 | 50 |
| LPS PB Ly6GInt | > | LPS SP Ly6GInt | 32 | |

**Table 2.** DEGs identified by p-value≤0.05 and fold change≥1.5 from comparisons of Ly6G$^{Int}$ neutrophil populations from the peripheral blood (PB), bone marrow (BM) and spleen (SP) of PBS-control and LPS-challenged mice. Data from transcriptomic analysis.

| Ly6G$^{Hi}$ PBS Comparisons: Blood, BM, SP, LV | | | DEGs | Total DEGs |
|---|---|---|---|---|
| PB Ly6GHi | > | BM Ly6GHi | 2274 | 3608 |
| BM Ly6GHi | > | PB Ly6GHi | 1334 | |
| PB Ly6GHi | > | SP Ly6GHi | 1628 | 1790 |
| SP Ly6GHi | > | PB Ly6GHi | 162 | |
| PB Ly6GHi | > | LV Ly6G+ | 1466 | 2106 |
| LV Ly6G+ | > | PB Ly6GHi | 640 | |
| BM Ly6GHi | > | SP Ly6GHi | 1882 | 3090 |
| SP Ly6GHi | > | BM Ly6GHi | 1208 | |
| BM Ly6GHi | > | LV Ly6G+ | 2652 | 5555 |
| LV Ly6GH+ | > | BM Ly6GHi | 2903 | |
| SP Ly6GHi | > | LV Ly6G+ | 1451 | 3733 |
| LV Ly6G+ | > | SP Ly6GHi | 2282 | |

**Table 3.** DEGs identified by p-value≤0.05 and fold change≥1.5 from comparisons of Ly6G$^{Hi}$ and Ly6G$^{+}$ neutrophils isolated from the peripheral blood (PB), bone marrow (BM), spleen (SP) and liver (LV) of PBS-control mice. Data from transcriptomic analysis.

| Ly6G$^{Hi}$ LPS Comparisons: Blood, BM, SP, LV | | | DEGs | Total DEGs |
|---|---|---|---|---|
| PB Ly6GHi | > | BM Ly6GHi | 591 | 1604 |
| BM Ly6GHi | > | PB Ly6GHi | 1013 | |
| PB Ly6GHi | > | SP Ly6GHi | 5 | 11 |
| SP Ly6GHi | > | PB Ly6GHi | 6 | |
| PB Ly6GHi | > | LV Ly6G+ | 108 | 323 |
| LV Ly6G+ | > | PB Ly6GHi | 215 | |
| BM Ly6GHi | > | SP Ly6GHi | 1153 | 1831 |
| SP Ly6GHi | > | BM Ly6GHi | 678 | |
| BM Ly6GHi | > | LV Ly6G+ | 2121 | 3236 |
| LV Ly6GH+ | > | BM Ly6GHi | 1115 | |
| SP Ly6GHi | > | LV Ly6G+ | 71 | 211 |
| LV Ly6G+ | > | SP Ly6GHi | 139 | |

**Table 4.** DEGs identified by p-value≤0.05 and fold change≥1.5 from comparisons of Ly6G$^{Hi}$ and Ly6G$^{+}$ neutrophils isolated from the peripheral blood, BM, spleen and liver of LPS-challenged mice. Data from transcriptomic analysis.

| PDAC Ly6G$^{Int}$: BM vs BM between genotypes | | | | | | |
|---|---|---|---|---|---|---|
| Geno | Pop | vs | Geno | Pop | DEGs | Total DEGs |
| WT | BMInt | > | KPflC | BMInt | 66 | 275 |
| WT | BMInt | < | KPflC | BMInt | 209 | |
| WT | BMInt | > | KPC | BMInt | 149 | 454 |
| WT | BMInt | < | KPC | BMInt | 305 | |
| KPflC | BMInt | > | KPC | BMInt | 1 | 1 |
| KPflC | BMInt | < | KPC | BMInt | 0 | |

**Table 5.** DEGs identified by p-value≤0.05 from comparisons of Ly6G$^{Int}$ neutrophils from the BM of WT, KP$^{fl}$C and KPC mice. Data from transcriptomic analysis. Genotype = Geno; population = Pop.

| PDAC Ly6G$^{Int}$: Spleen vs Spleen between genotypes | | | | | | |
|---|---|---|---|---|---|---|
| Geno | Pop | vs | Geno | Pop | DEGs | Total DEGs |
| WT | SPInt | > | KPflC | SPInt | 401 | 707 |
| WT | SPInt | < | KPflC | SPInt | 306 | |
| WT | SPInt | > | KPC | SPInt | 865 | 1419 |
| WT | SPInt | < | KPC | SPInt | 554 | |
| KPflC | SPInt | > | KPC | SPInt | 0 | 0 |
| KPflC | SPInt | < | KPC | SPInt | 0 | |

**Table 6.** DEGs identified by p-value≤0.05 from comparisons of Ly6G$^{Int}$ neutrophils from the spleen of WT, KP$^{fl}$C and KPC mice. Data from transcriptomic analysis. Genotype = Geno; population = Pop.

| PDAC Ly6G$^{Int}$: Unique and Shared DEGs compared to WT | | | | | | |
|---|---|---|---|---|---|---|
| Tissue | Geno | Comparison | | | | Total DEGs |
| BM | WT | BMInt | > | KPflC & KPC | 40 | 187 |
| | | BMInt | < | KPflC & KPC | 147 | |
| | WT | BMInt | > | KPflC Only | 26 | |
| | | BMInt | < | KPflC Only | 62 | 88 |
| | WT | BMInt | > | KPC Only | 109 | |
| | | BMInt | < | KPC Only | 158 | 267 |
| Spleen | WT | SpleenInt | > | KPflC & KPC | 290 | 504 |
| | | SpleenInt | < | KPflC & KPC | 214 | |
| | WT | SpleenInt | > | KPflC Only | 111 | 204 |
| | | SpleenInt | < | KPflC Only | 93 | |
| | WT | SpleenInt | > | KPC Only | 575 | 916 |
| | | SpleenInt | < | KPC Only | 341 | |

**Table 7.** DEGs identified by p-value≤0.05 from the BM and spleen that are shared by KP$^{fl}$C and KPC compared to WT, only identified in WT vs KP$^{fl}$C and only identified in WT vs KPC mice. Data from transcriptomic analysis. Genotype = Geno.

| PDAC Ly6G$^{Int}$: BM vs Spleen within each genotype | | | | | |
|---|---|---|---|---|---|
| Geno | Comparison | | | DEGs | Total DEGs |
| WT | BMInt | > | SPInt | 461 | 1135 |
| | BMInt | < | SPInt | 674 | |
| KPflC | BMInt | > | SPInt | 6 | 155 |
| | BMInt | < | SPInt | 149 | |
| KPC | BMInt | > | SPInt | 12 | 228 |
| | BMInt | < | SPInt | 216 | |

**Table 8.** DEGs identified by p-value≤0.05 from comparisons of Ly6G$^{Int}$ neutrophils from the BM and spleen in WT, KP$^{fl}$C and KPC mice. Data from transcriptomic analysis. Genotype = Geno.
